# Antibody-recruiting molecules based on mimotope fusion peptides and a dimeric antibiotic

**DOI:** 10.64898/2026.09.29.755485

**Authors:** Franz G. Zingl, Meilee L. Mekalanos, Deborah R. Leitner, Stephen R. Walsh, Matthew K. Waldor, John J. Mekalanos

## Abstract

New therapeutic approaches are needed to treat infections due to multidrug resistant (MDR) Gram-negative bacteria. The use of antibody-recruiting molecules (ARMs) is a promising approach to address the MDR problem given their potential to engage both the adaptive and innate immune systems to treat bacterial infections. Here we present two new strategies to create ARMs. First, we show that a 9-mer antimicrobial peptide (K6) can be fused to a 11-mer mimotope peptide recognized by antibodies targeting the *Vibrio cholerae* O1 O-antigen to produce an ARM peptide (FZ5) that can render *E. coli* more sensitive to complement killing in the presence of antibodies that recognize this mimotope. FZ5 can drive killing by complement in the presence of rabbit antibodies raised against O1 serogroup *V. cholerae* as well as antibodies present in human sera elicited by immunization with the live attenuated cholera vaccine PanChol. We also describe a strategy that combines bacterial surface binding with hapten display through the use of homodimers of the antibiotic streptomycin (Str) and anti-Str antibodies elicited by Str-protein conjugate vaccines. When combined with dimeric Str derivatives and human complement, polyclonal anti-streptomycin antisera kills *E. coli* by targeting the Str hapten. Dimeric forms of Str also significantly enhanced killing of a Str-resistant *P. aeruginosa* clinical isolate dependent on low levels of human complement and polyclonal anti-Str antibody. Together these data suggest that anti-bacterial ARM molecules based on mimotope display through simple synthetic fusion peptides, or dimeric haptens have promise as new approaches for the treatment of MDR bacterial infections.

## INTRODUCTION

Multidrug-resistant (MDR), Gram-negative pathogens are an increasing cause of life-threatening infections and account for at least 700,000 deaths every year (1). These pathogens include species such as *Pseudomonas aeruginosa*, *Escherichia coli, Klebsiella pneumoniae*, and *Acinetobacter baumannii* (2). Infections due to these increasingly resistant bacteria can require treatment with antibiotics that have unfavorable side effect profiles such as polymyxins (3). The emergence of MDR Gram-negative bacteria is a particularly worrisome threat because so few novel drugs are in the development pipeline that can address this problem. Clearly, fundamentally new approaches are needed to address the treatment of infections due to MDR Gram-negative organisms that are less prone to rapid evolved resistance (4,5).

Antibody-recruiting molecules (ARMs) represent a promising approach to address the MDR problem by engaging elements of both the adaptive and innate immune systems (6). Pioneering work over the last decade has laid the foundation for the ARM concept as a therapeutic modality for engaging complement-dependent cytotoxicity (CDC), antibody-dependent cellular cytotoxicity (ADCC), and opsonization as mechanisms for targeted killing of bacteria *in vitro* or during infection (7). In these studies, a bifunctional ARM composed of a bacterial cell binding motif fused to a small molecule hapten was used. Typically hapten molecules such as 2,4-dinitrobenzene, α-galactose, or L-rhamnose were selected for ARM designs because antibodies against these haptens exist in normal human serum and these molecules are thought to be absent from mammalian host cell surfaces (8). However, the affinity of such ‘natural antibodies’ for these haptens is generally very low (9) probably because the antibodies are produced by germline genes and likely did not arise by B-cell hypermutation and affinity maturation. ARM approaches that take advantage of active immunization with hapten-bearing or cross-reacting immunogens is not apparent in the literature.

The molecules utilized by reported ARMs to decorate microbial cells with haptens also vary. They include specific monosaccharides that have affinity for bacterial receptors (11), antibiotics such as polymyxin and vancomycin that bind bacterial cell envelope components (12), or siderophore-iron complexes which have high affinity for cognate receptors involved in bacterial iron scavenging (13). However, in general the chemistry required to assemble these types of ARM therapeutics is challenging and not readiy available to most microbiology laboratories. We have previously explored the development of antimicrobial peptides (AMPs) that have activity against Gram-negative cells and reported that Peptide K6 has promising activity against important Gram-negative species (14). K6 is a symmetrical, linear 9-mer peptide with the composition WWKKWKKWW (single letter residue W= tryptophan, K= lysine); the K6 amino terminus is not blocked and its caboxy terminus is amidated (14). Thus, K6 is predicted to have a net charge of +5 at physiological pH and its cationic nature suggests that it likely binds the lipid A moiety of lipopolysaccharide which is displayed on the surface of Gram-negative cells similar to what is postulated for polymyxins and many other AMPs (18). While the mechanism of action of K6 is not well-defined, its ability to form nanomicelles suggested that K6 could be used to assemble dense, multimeric arrays of K6-linked epitopes on the surface of Gram-negative cells. Mimotopes are typically peptides that can react with antibodies directed against antigens of diverse chemical composition (15). We therefore speculated that mimotope peptides fused to K6 would be directed to the surface of organisms such *E. coli* where binding of anti-mimotope antibodies would trigger antibody-dependent cell killing in the presence of complement. Thus, we hypothesized that ARMs based on AMPs fused to peptide mimotopes represented an opportunity to develop a new class of ARMs where simple peptide chemistry could be utilized in their synthesis.

Here, we report that K6 is able to display a mimotope peptide on the surface of *E. coli* cells that renders them more sensitive to complement killing in the presence of antibody targeting the mimotope. The mimotope we selected (A12X) is recognized by anti-O1 serogroup *Vibrio cholerae* rabbit antisera (16) and apparently human sera elicited by immunization with the live attenuated cholera vaccine PanChol (17). Our initial positive results with A12X displayed by K6 prompted us to design a new strategy that combines both bacterial surface binding with hapten display. This approach involved development of homodimers of the antibiotic streptomycin (Str) and Str-protein conjugate immunogens that elicited anti-Str antibody responses. These polyclonal anti-Str hapten sera led to killing of clinical isolates of *E. coli* and *P. aeruginosa* (18–20). Our results suggest that the modified approaches to ARM development described here warrant further investigation for control of Gram-negative pathogen infection.

## RESULTS

### Testing of K6 peptide fusions designed to display mimotopes of the *Vibrio cholerae* O1-antigen for their ability to target complement-dependent killing of *E. coli*

Mimotopes composed of synthetic peptides have been identified that imitate epitopes present on protein or carbohydrates antigens (15). They have typically been identified by phage-display of short, random, peptides which bind to monoclonal or polyclonal antibodies directed against the antigen of interest. The *V. cholerae* O1-antigen represents the principal target of protective antibodies against the cholera pathogen and consists of a homopolymer of acylated perosamine; the two distinct O1 serotypes, termed Ogawa and Inaba, only vary based on the methylation of a terminal perosamine residue (21, 22). Mimotopes of the *V. cholerae* O1-antigen have been reported in the literature and were identified by phage-display using either monoclonal or polyclonal anti-O1 antibodies (16, 23–25). We selected five of these published O1 mimotope peptides and fused them to the C-terminus of K6 separated by a flexible spacer tripeptide GPG to produce candidate ARM fusion peptides (**Fig. 1, left panel**). These synthetic peptides, which range in length from 22 to 32 residues in length, were produced by a commercial vender (GeneScript) at greater than 96% purity and thus represent potential ARMs based entirely on synthetic peptides with no coupling chemistry required.

**Fig. 1.**
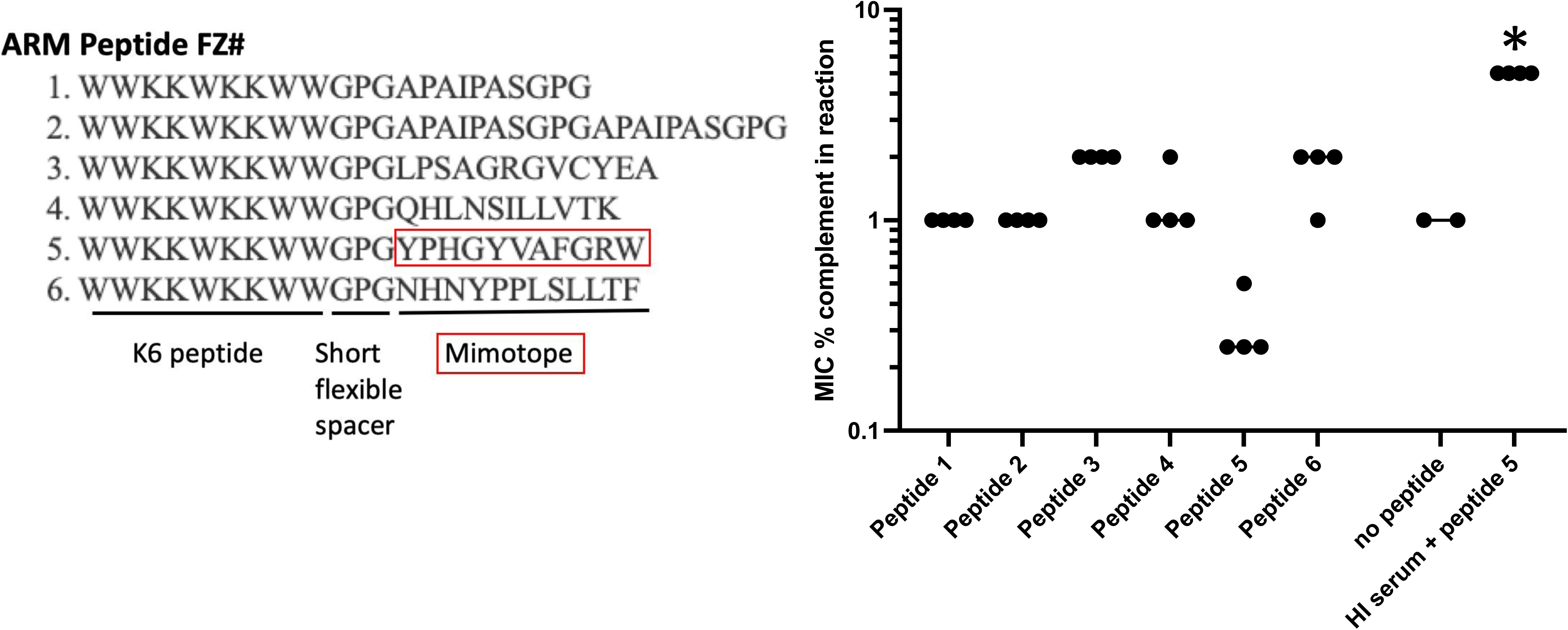
ARM peptide fusions between peptide K6 and *V. cholerae* O1 antigen mimotope peptides and their ability drive complement-dependent and anti-O1 serum bactericidal activity. Left. Sequence of ARM peptides FZ1-FZ6 in standard single letter amino acid code from N-terminus to C-terminus. The location of K6 sequence is indicated as is the flexible linker sequence (GPG) separating it from the mimotope sequences. An example of the published (16) mimotope A12x is designated with the red boxes. **Right.** Sensitivity *of E. coli* OmniMAX promoted by the indicated ARM peptides 1-6 compared to controls expressed as the minimum inhibitory concentration (MIC) of human complement (%) that produced total killing of the bacteria present in the assay. Bacteria were exposed to the ARM peptides at approximately the concentration of 50 μM before exposure to complement for 1 hr. The asterisk denotes that heat inactivated (HI) human complement was not tested above ∼5% final concentration in the assembled reaction.

*E. coli* K12 strain OmniMAX cells were coated with each of the six synthetic ARM peptides and then exposed to commercial anti-O1 rabbit typing sera (Difco) in the presence of various concentrations of human complement. As shown in **Fig. 1 (right panel),** one peptide called FZ5 caused hypersensitivity to human complement in the presence of anti-O1 sera. Heat inactivation of the complement at 56°C for 30 mins eliminated this killing effect. This enhanced bacteriocidal activity of human complement was not observed in the absence of the FZ5 peptide.

### *E. coli* OmniMAX and and mutants defective in PNAG production are highly sensitive to human complement

Further analysis of this bacteriocidal effect associated with FZ5 combined with anti-O1 rabbit sera revealed that the strain OmniMAX was highly sensitive to human complement obtained from Innovative Reseach even in the absence of anti-O1 antibodies (**Fig 2, left panel**). Accordingly, we expanded our studies to E11, a K1 encapsulated *E. coli* isolated from a neonatal sepsis case (25, 26). Besides the K1 capsular polysaccharide, this strain also produces a polysaccharide called poly-B-(1–6)-N-acetyl-D-glucosamine (PNAG) (27). Because PNAG is a common antigen produced by many bacterial species we elected to use a mutant of E11 defective in PNAG production (E11Δpga) for our initial studies (25, 26) in order to avoid anti-PNAG antibody-dependent killing that might occur if the rabbit or human sera providing complement contained such anti-NAG antibodies. Spontaneous streptomycin (Str) resistant mutants of E11 and E11Δpga were generated, E11 StrR and E11Δpga StrR, respectively in order to use these same strains for studies of Str-based ARM molecules (see below).

**Fig. 2.**
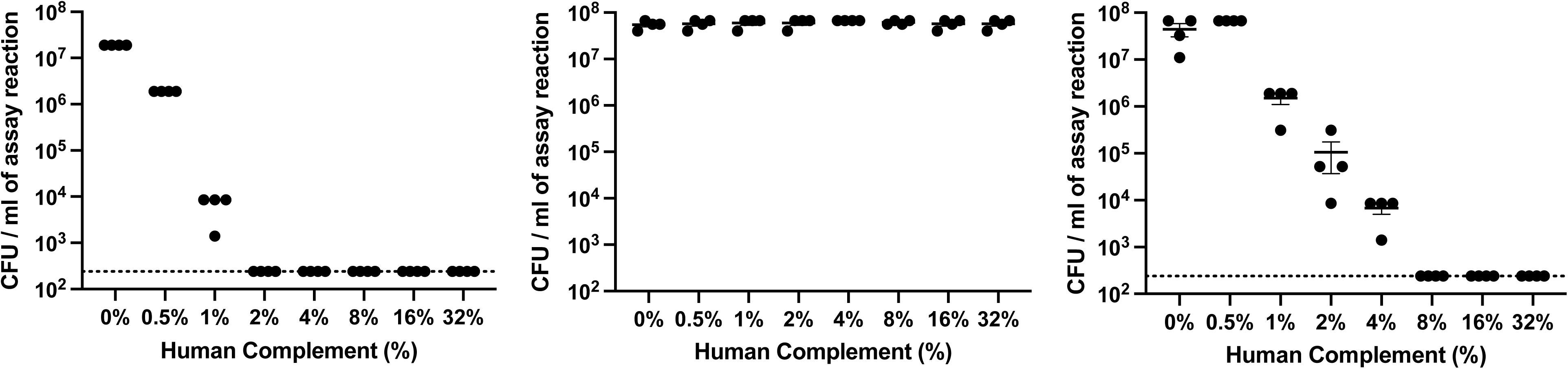
Comparison of *E. coli* OmniMAX, E11 and its mutant defective in PNAG production for sensitivity to human complement. Left. Sensitivity of *E. coli* OmniMAX to human complement concentrations indicated in a standard 1 hr assay performed in PBS. **Middle.** Sensitivity of an *E. coli* E11 mutant defective in PNAG production (E11Δpga StrR) to human complement. **Right.** Sensitivity PNAG^+^ E11 StrR to human complement. Concentrations of human complement are indicated in a standard 1 hr assay performed in PBS.

As shown in **Fig. 2 (right panel)**, we found that the PNAG-defective mutant E11Δpga StrR was at least 30-fold more sensitive to Innovative Research complement compared to the PNAG^+^ strain E11 StrR (**Fig. 2, middle panel**). The streptomycin-sensitive parent (E11Δpga) of E11Δpga StrR also displayed hypersensitivity to this source of human complement (data not shown). This bacteriocidal activity was heat-labile when treated at 56°C for 30 mins suggesting it was associated with complement components. We did not explore whether the absence of PNAG expression in E11Δpga StrR was responsible for the apparent increased complement sensitivity observed by a more rigorous analysis but rather elected to simply use this hypersensitive strain for extended studies on antibody-dependent complement killing in the presence of the ARM molecules we were designing (see below).

### Anti-O1 rabbit serum can drive enhanced killing of *E. coli* E11Δpga StrR in the presence of human complement and FZ5

We next tested if the FZ7 ARM peptide could drive killing of E11Δpga StrR in the presence or absence of human complement and anti-O1 rabbit typing sera. As shown in **Fig. 3 (left panel),** Peptide FZ5 alone (at 2 μg/ml in the assay) sensitized E11Δpga StrR to killing by 0.5% human complement in the absence of anti-O1 antisera. Addition of anti-O1 rabbit typing sera to 1% in the presence of human complement produced another order of magnitude of killing of E11Δpga StrR that reached statistical significance compared to complement alone, or complement plus antisera in the absence of peptide FZ5. Importantly, peptide K6 (i.e., the cell-binding, targeting AMP for FZ5) alone or in combination with anti-O1 serum, with or without human complement, did not enhance killing of E11Δpga StrR (**Fig 3, right panel**), suggesting that the enhanced FZ5-dependent killing cannot be explained by the properties of K6 alone. Other assays showed that the MIC of FZ5 was in excess of 32 μg/ml in Luria broth for strain E11Δpga StrR (data not shown) indicating that the bacteriocidal activity observed for FZ5 in the presence of anti-O1 rabbit antisera and human complement was likely complement and anti-O1 antibody dependent. Together these data suggest that the FZ5 ARM peptide can bind to *E. coli* E11Δpga StrR cells through its K6 moiety and trigger enhanced killing of these bacterial cells by displaying the A12X O1 mimotope to anti-O1 antibodies and human complement.

**Fig. 3.**
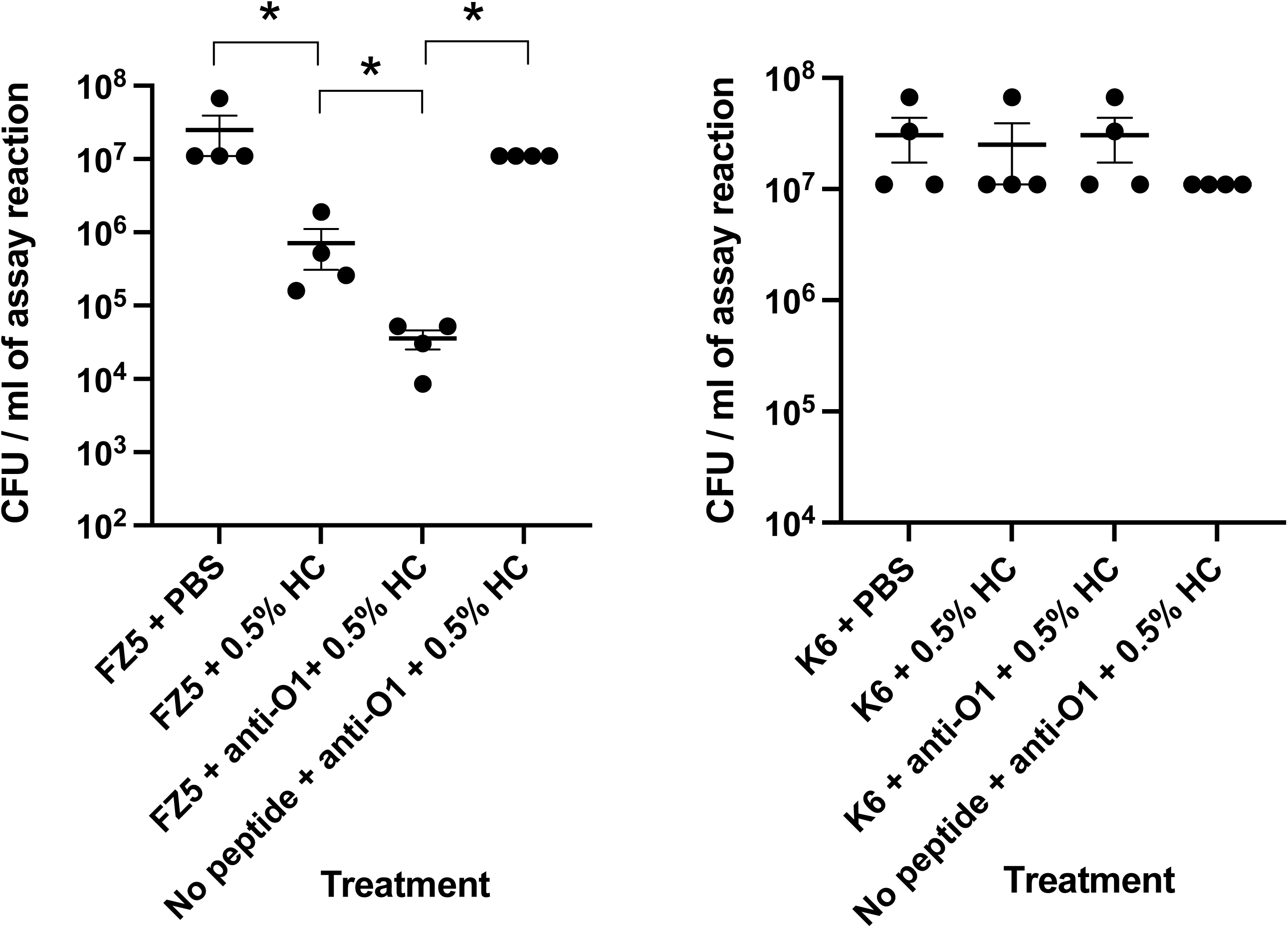
Sensitivity of *E. coli* to peptides FZ5 or K6 with or without *V. cholerae* rabbit anti-O1 typing antisera in the presence or absence of human complement (HC). Assays were performed on E11Δpga StrR and viable counts (CFU) per ml of assay are shown after 1 hr incubation at 37°C. **Left.** Bactericidal activity of the FZ5 peptide (at a final concentration 2 μg/ml) in the presence or absence of 1% anti-O1 rabbit antisera with or without human complement (HC)(0.5%). Anti-01 serum plus HC was also tested in the absence of FZ5. **Right.** Bactericidal activity of the K6 peptide (at a final concentration 2 μg/ml) in the presence or absence of 1% anti-O1 rabbit antisera with or without HC (0.5%). Anti-01 serum plus HC was also tested in the absence of FZ5. * Mann-Whitney test, value < 0.05 for indicated comparison data sets (brackets).

### Human sera from volunteers immunized with oral cholera vaccine PanChol promotes killing of *E. coli* in the presence of FZ5 and complement

PanChol is a promising live attenuated *V. cholerae* vaccine capable of inducing strong anti-O1 immune responses in human subjects after a single oral dose (17). We tested if sera from volunteers immunized with this vaccine could also kill *E. coli* E11Δpga StrR in the presence of the O1 mimotope FZ5 ARM peptide and human complement. Pools of heat-inactivated, pre-immunization (Day 1) and post-immunization (Day 15) sera were prepared for these experiments from PanChol vaccinees. Peak vibriocidal titers are typically observed at Day 15 showing ∼1500-fold increase in geometeric mean titer indicating that PanChol elicits a potent anti-*V. cholerae* antibody response that likely consists of IgM and IgG antibodies directed against the O1 antigen (17). Accordingly, we tested these two human sera pools for antibodies (after heat inactivation at 56°C for 30 mins) for their ability to kill *E. coli* E11Δpga StrR when mixed with FZ5 peptide in the presence of human complement.

As shown in **Fig. 4** the combination of FZ5 at 2μg/ml and 1/100 dilution of serum from the Day 15 post-PanChol vaccinees caused approximately a 10-fold decrease in the recovery of viable cells compared to the same dilution of Day 1 sera or compared to reactions that contained no sera other than 0.5% human complement. This killing effect also depended on the presence of FZ5 peptide, and heat-inactivation of the human complement eliminated the bactericidal effect observed. We found that coating with K6 alone did not promote enhanced killing of E11Δpga StrR cells in the presence of Day 15 post-PanChol sera (**Fig. 4**). Together, these studies suggest that active immunization of human subjects with PanChol can induce antibodies that recognize the mimotope displayed by ARM peptide FZ5. Although the bactericidal effects observed were relatively modest in magnitude (about 10-fold enhanced killing compared to controls), these results suggest that FZ5 could display a peptide mimotope on the surface of bacterial cells using its cationic peptide K6 moiety to target the Gram-negative cell evelope and that this presentation of the mimotope, in turn, could enable the killing of cells with vaccine-elicited antibodies and human complement.

**Fig. 4.**
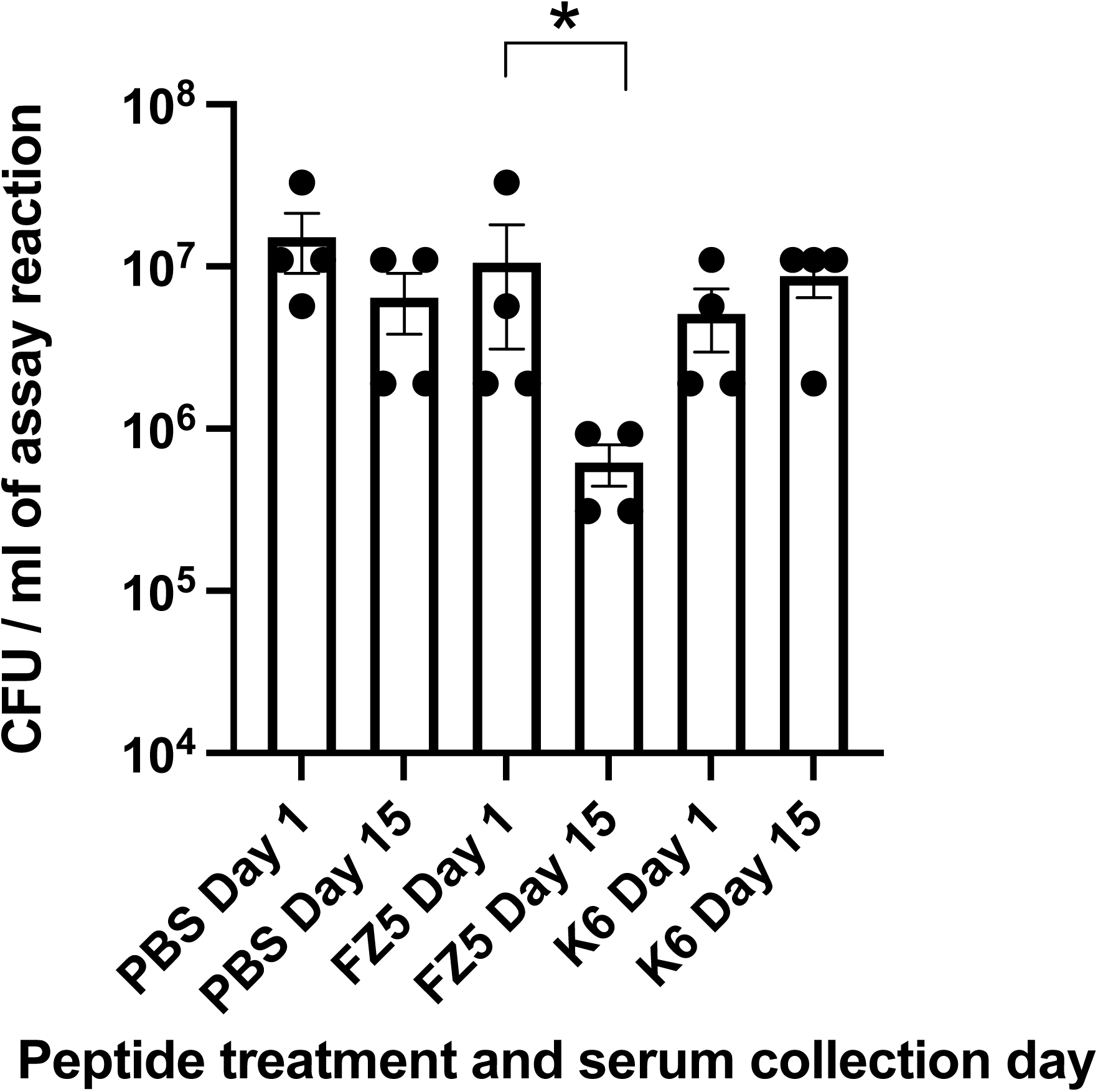
Sensitivity of *E. coli* to peptide FZ5 or K6 with or without *V. cholerae* anti-O1 antisera generated by PanChol immunization. Assays were performed on E11Δpga and viable counts (CFU) per ml of assay are shown after 1 hr incubation at 37°C in PBS. Peptides FZ5 or K6 were added where indicated at a final concentration 2 μg/ml. The indicated pooled sera from the same set of volunteers were collected at Day 1 or Day 15 after oral immunization with PanChol as described previously (17) and were heat inactivated at 56°C for 30 mins and then mixed to be at a final concentration of 1% in the assays as indicated. Human complement was added to each assay at a final concentration of 0.5% as the last step in the assembly of the reactions. * Mann-Whitney test, value < 0.05 for indicated comparison data sets (brackets).

### Development of modified monomeric and dimeric streptomycin derivatives

FZ5 is 23 L-amino acid residues in length and could have poor pharmacological properties because of its large molecular size and likely suceptibility to proteolysis. We wondered if cell binding and epitope display could be combined in one non-peptide small symmetrical molecule that was potentially simple to synthesize and highly stable. We have named such a ARM-related compounds ‘<u>H</u>omodimeric <u>A</u>ntibody <u>R</u>ecruiting Molecules’ or ‘HARMs’ to differentiate this concept from ARM molecules described in the literature which are always heterodimers (6–8). Furthermore, because there is evidence that some antibiotics assemble into multimeric complexes on bacterial surface (28, 29), we hypothesized that such multimeric complexes could enhance recruitment of antibody molecules in the density needed to trigger ADCC and opsonization activity (30).

We selected streptomycin (Str) to explore the HARM concept for several reasons. While dimers and heterodimers of aminoglycosides have previously been reported (31) these molecules were designed to maintain their inherent antibiotic mechanism. In our study design, we intended to use Str-resistant target strains to eliminate the normal mechanism of action of this antibiotic. Str is a cationic molecule with a +3 charge under physiological pH, small in size (only 582.27 Da as a +1 ion), highly stable *in vivo*, and further displays a unique aldehyde group that is available for simple chemical modifications through reductive amidation (see below). Str is seldom used as a antibiotic in humans for treatment of Gram-negative infections because of various adverse events associated with its use at effective therapeutic levels (32). Thus, we speculated that if antibodies against the drug could improve its efficacy *in vivo* by, for example, by engaging complement-dependent killing mechanisms, this might allow much lower doses of Str-based HARM analogs to be used to treat infections. Commercially available anti-Str monoclonal antibodies (33) also suggested that Str is a reasonable hapten to explore in such a HARM application. Accordingly, we developed simple Str-carrier protein conjugate immunogens (34) for raising polyclonal anti-Str antibodies. An overview of the program we developed follows.

### Reductive methylation in the presence of organic diamines as a synthetic strategy to make dimeric forms of streptomycin

We reasoned that a dimeric form of Str might be less prone to enter bacterial cells because of its size but still retain or even enhance its surface binding properties due to its cationic nature. Such dimers could display a Str epitope at either of their ends in solution phase to allow the Str eptitope to engage anti-Str antibodies on its apical end while still binding target bacterial cells on its distal end.

We chose three linkers for the initial investigation of this concept – 1) Ethylene diamine (Eda), a small two carbon linker that we planned to use primarily for making monomeric Str derivatives; 2) Diethylene glycol bis (3-amino propyl) ether (LinkB) a longer hydrophilic linker; and 3) Cystamine (Csa), a molecule related to the amino acid cysteine. We predicted that these linkers would allow stable (Eda and LinkB) or cleavable (Csa) dimers to form. When added to a monomer of Str, all three linkers would provide a unique primary amino group for coupling to carrier proteins for use as conjugate Str-protein immunogens. The details of our synthetic strategies are provided in METHODS.

In brief, synthesis involved aqueous reductive amidation of the three linkers with Str in separate reactions using sodium borohydride as a reducing agent (**Fig. 5**). Acetone precipitation of Str derivatives, extraction of water soluble products, and then extensive dialysis (using <1000 Da cut-off membranes) followed by lyophilization was used to produce final products used in the analysis. Liquid chromatography coupled to mass spectroscopy (LCMS) provided analytical evidence that Str molecules were quantitatively reduced to yield multiple expected products in each of the synthetic reactions (**Fig. 6**). However, the ratios of linker to Str in the designed reactions drove production of primarily monomeric modified Str products (i.e., Str-Eda reaction) or dimeric products (Str-LinkB and Str-Csa reactions). We used the ratio of monomeric Str derivatives to dimeric mass derivatives to predict the likely abundance of dimeric species and made no attempt to separate monomeric from dimeric species in these reactions for our analysis of their potential activities.

**Fig. 5.**
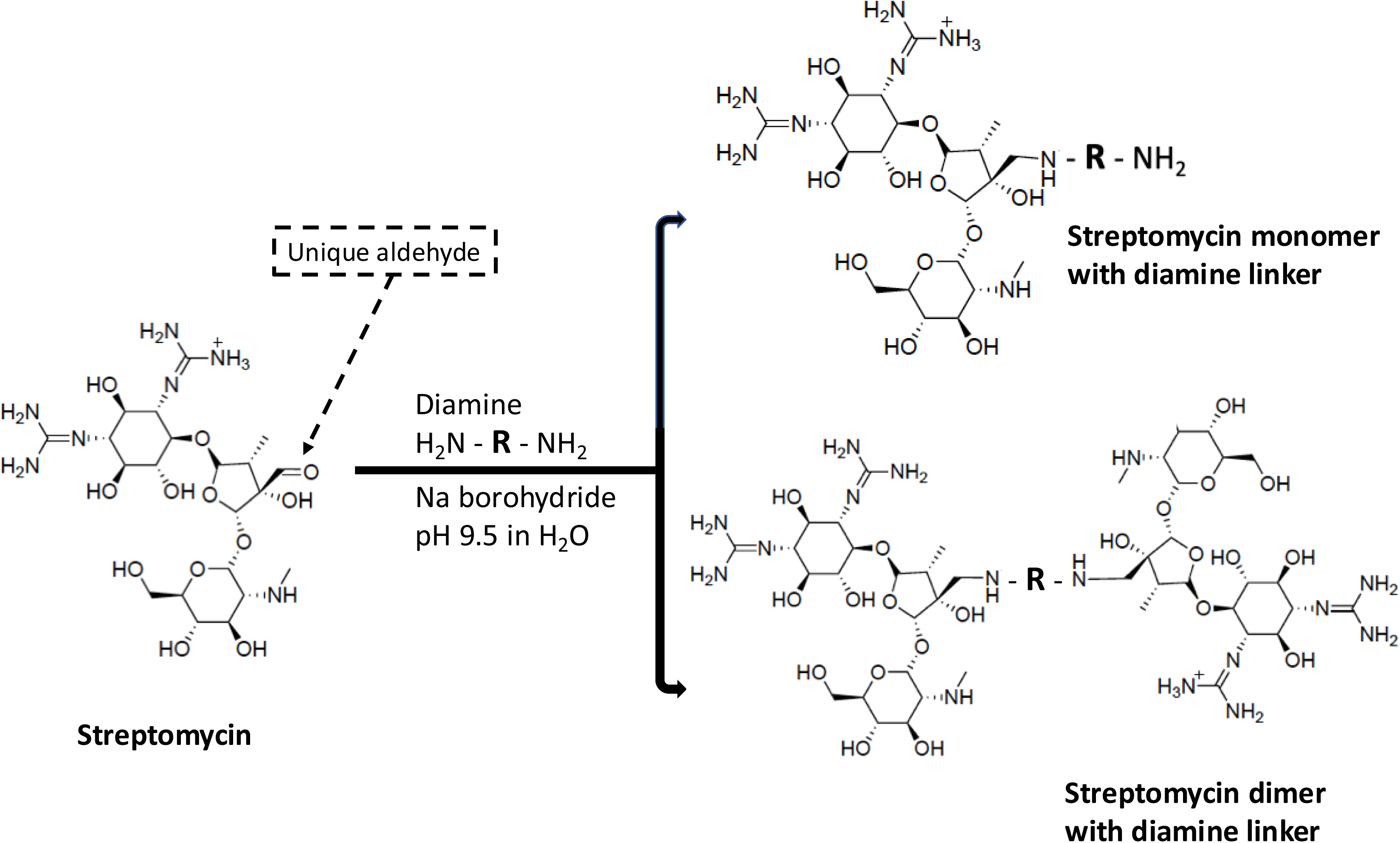
Schematic diagram of reductive amidation reaction of streptomycin with diamines to produce monomer and dimer molecules. The single aldehyde group in streptomycin (dashed arrow) was reacted with various diamines (see text and METHODS for details) to form Schiff bases which were then reduced with sodium borohydride. This synthetic strategy produced a mixture of products including mono-substituted secondary amines (top right), streptomycin dimers (bottom right) and reduced di-hydro-streptomycin)(not shown). Examples of these three classes of products are shown in Fig. 7 and 8.

**Fig. 6.**
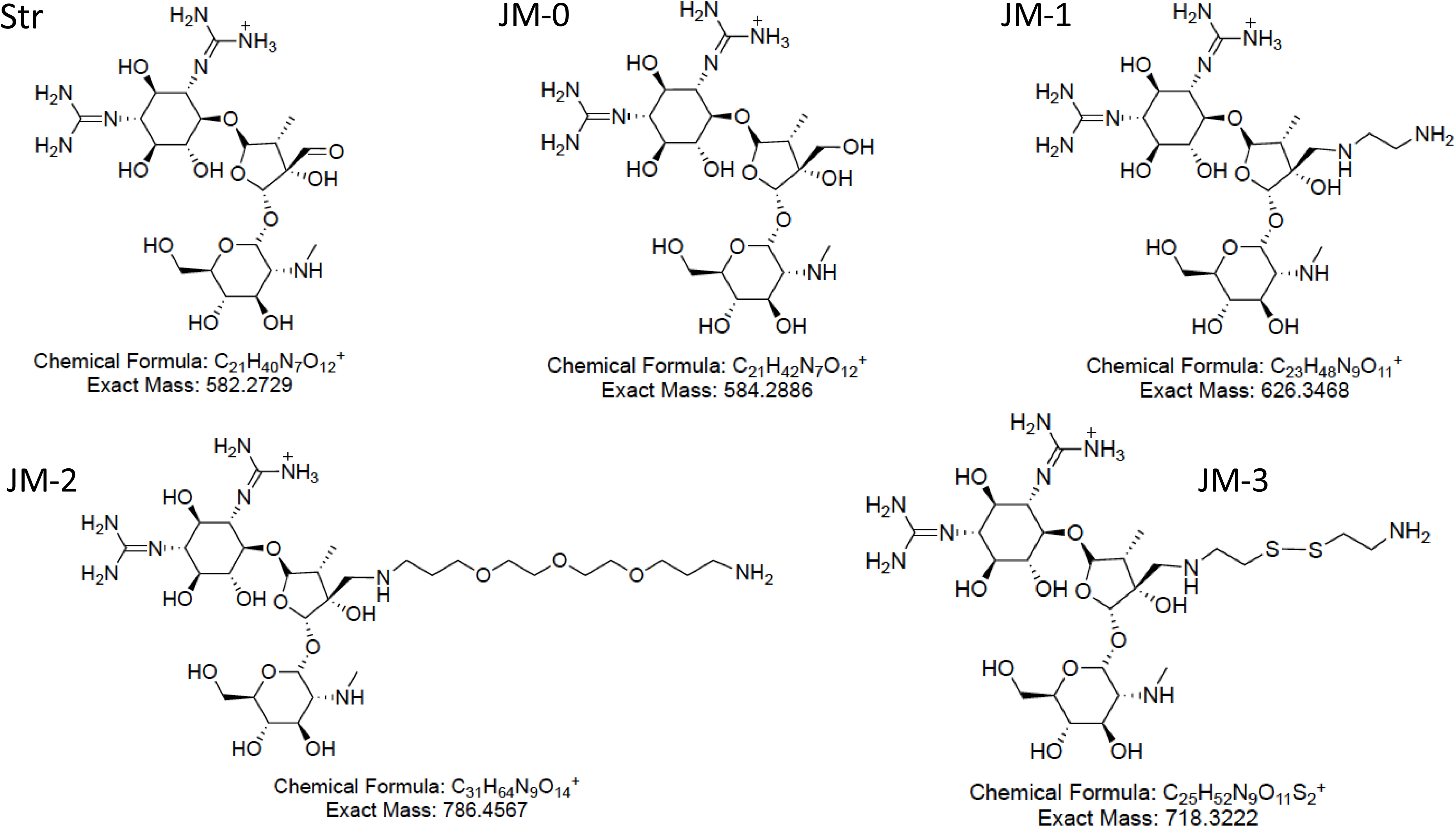
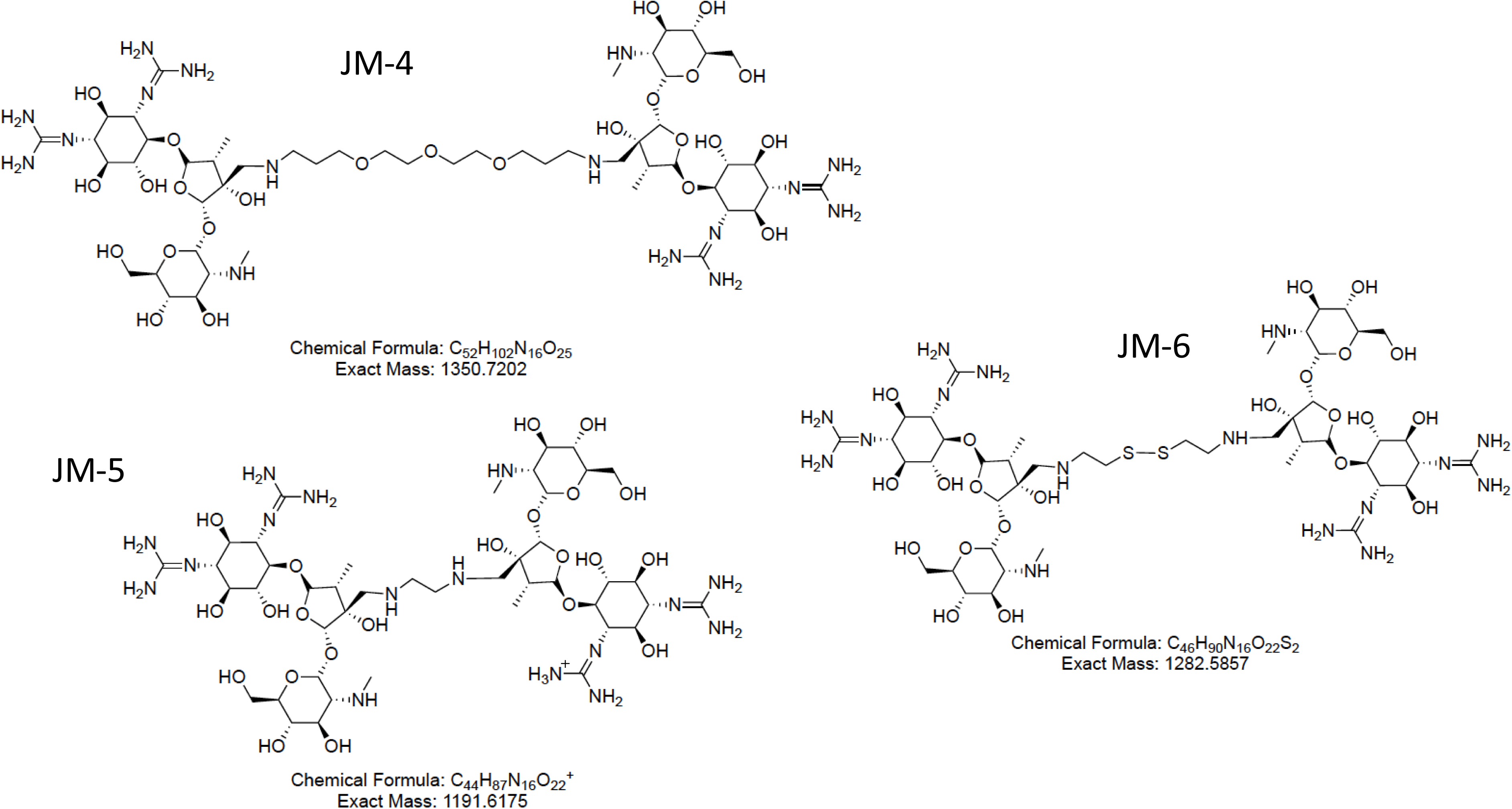
Expected structures of precursors and products of reductive amidation of streptomycin in the presence of the three diamines used in our study. Top. Streptomycin once reduced is expected to generate di-hydro-streptomycin (JM-0) in all reactions. The mono substituted products of reaction with ethylene diamine (Eda)(JM-1), diethylene glycol bis (3-amino propyl) ether (LinkB)(JM-2) or cystamine (Csa)(JM-3) are shown in the top half of the figure. **Bottom.** Shown are the expected streptomycin dimer products of LinkB (JM-4), Eda (JM-5) and Csa (JM-6). The predicted chemical formulas and exact molecular masses of these chemical species are shown below their structures.

#### Str-Eda

We anticipated that because of the ratio of Eda to Str used in the reductive methylation reaction (Eda 50-molar excess), the monomeric Str adduct was the expected product. The LC profile revealed a single homogeneous peak that was composed of mass species corresponding to monomeric Str-Eda (626.35 Da as the +1 ion)(JM-1) with a small amount of reduced Str (584.29 Da as a +1 ion)(ML-0)(**Fig. 7, Top**). Very little dimeric (Str)_2_ -Eda (predicted 1191.6 Da for +1 detected)(JM-5) was detected. Based on these findings, we selected the Str-Eda reaction products to prepare conjugate immunogens for generating anti-Str polyclonal antibodies (see below).

**Fig. 7.**
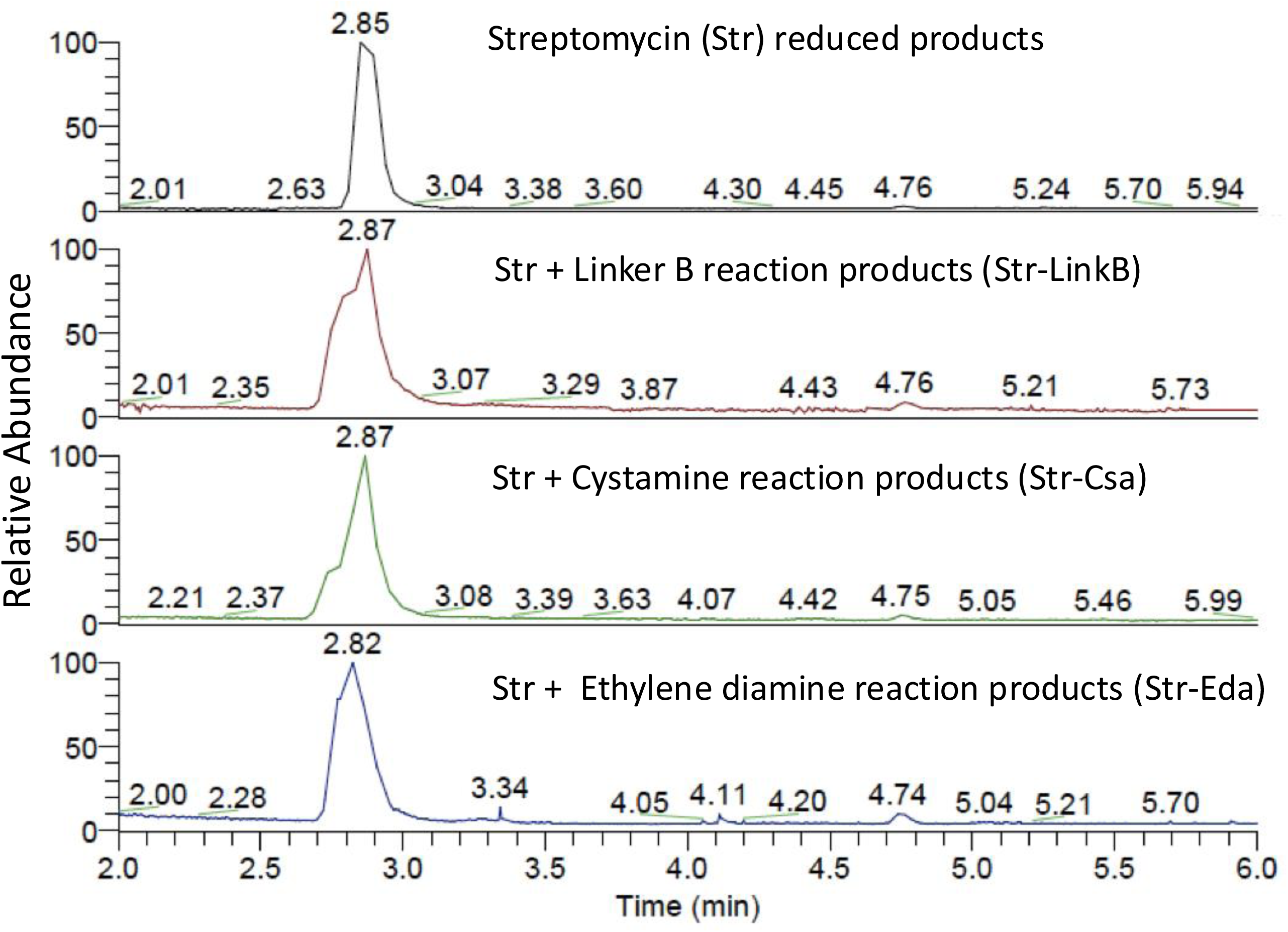
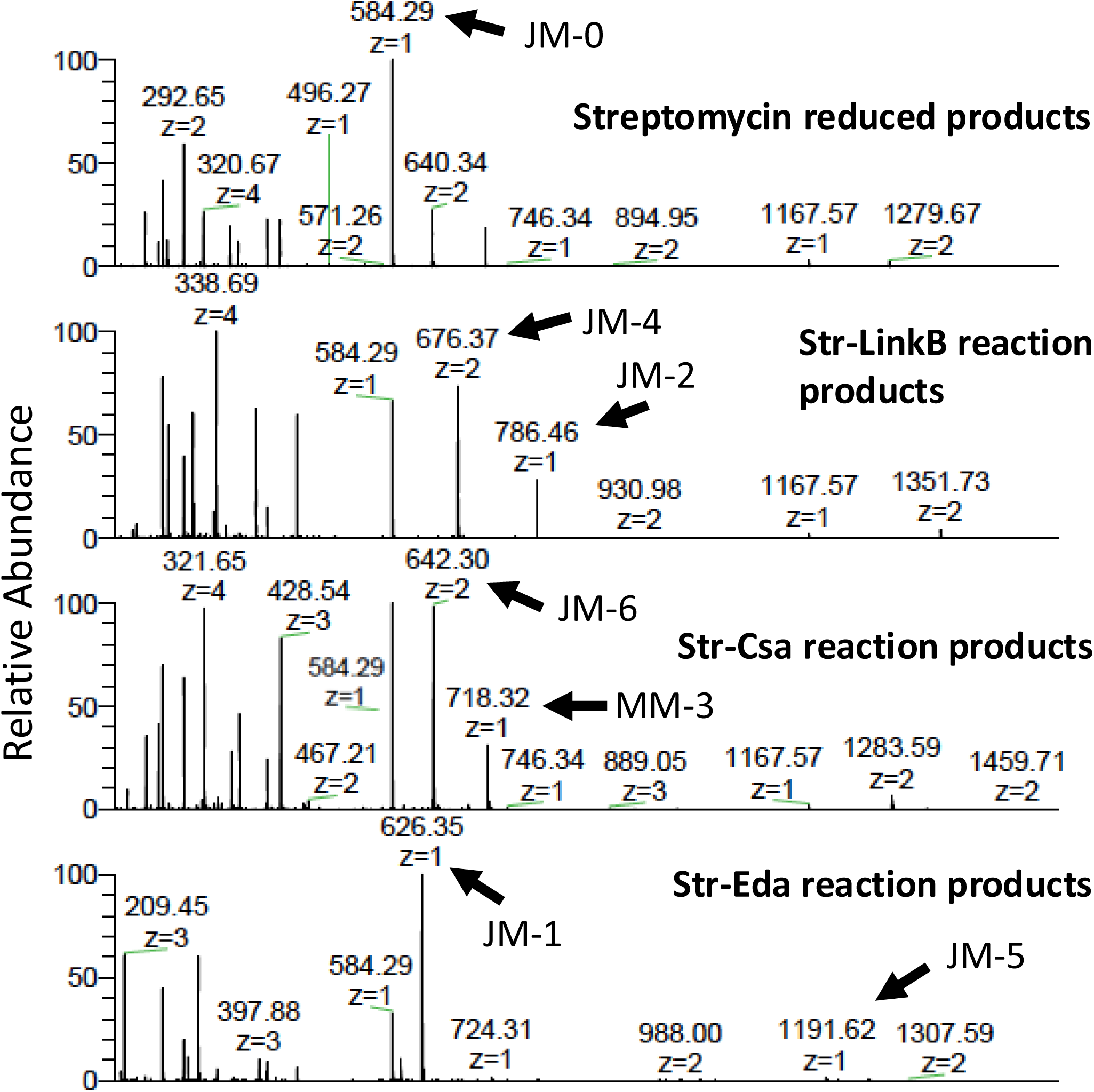
Elution times and profiles of products from sodium borohydride reactions of streptomycin alone or with the three diamines used in our study. Temporal elution profile of products from a C18 column loaded with the Str-Eda, Str-LinkB and Str-Csa reaction products (**Top**) before analysis by tandem mass spectrometry. Relative absorbance mass species (**Bottom**) resolved by the mass spectrometer operating in positive-ion mode.

#### Str-LinkB

The ratio of LinkB to Str (20:1 ratio) reactions designed predicted more dimeric adduct would form and our analysis confirmed this. A bifurcated peak emerged form LC (**Fig. 7, top**) and mass species in this elution peak corresponding to monomeric Str-LinkB (786.46 Da as the +1 ion)(JM-2) with nearly an equal amount of a the predicted dimer (1350.72 as the +2 ion)(JM-4); reduced or dihydro Str (584.29 Da as a +1 ion)(JM-0) was also present in the Str-LinkB reaction (**Fig. 8, bottom**).

**Fig. 8.**
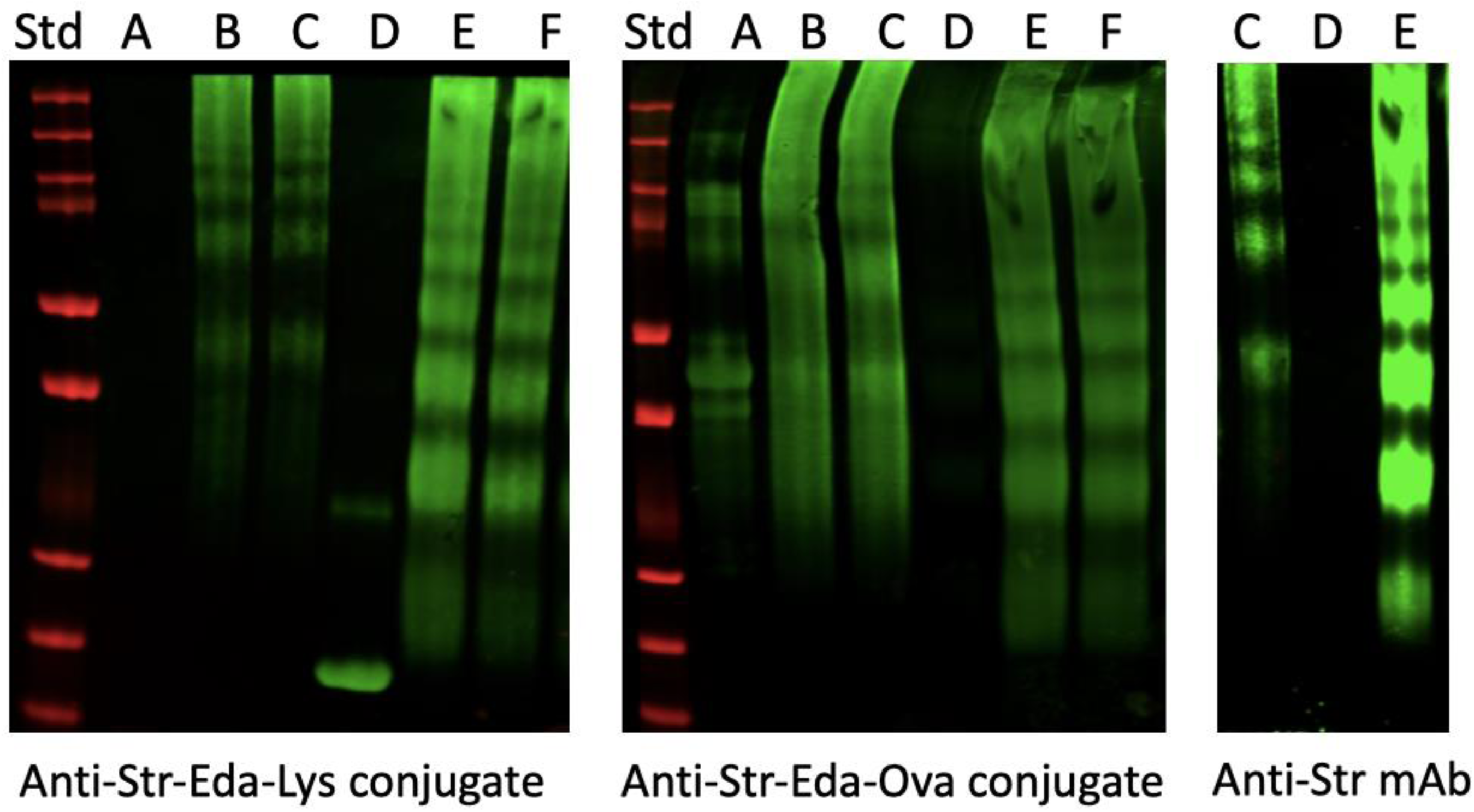
Detection of Str haptins by western blot analysis. Carrier proteins ovalbumin (Ova) and hen egg white lysozyme (Lys) were coupled to the Str-Eda derivatives in the presence of glutaraldehyde. These were the same immunogens used to generate rabbit anti-Str-Eda-carrier protein rabbit sera used in the analysis. Samples of the conjugates were subjected to polyacrylamide gel electrophoresis in the presence of sodium dodecyl sulfate, transferred to membranes, probed with the indicated antisera under each panel, followed by appropriate secondary antibodies coupled to the enzyme horse radish peroxidase (HRP). Anti-Str-Eda-Lys (**Left**) and Anti-Str-Eda-Ova (**Middle**) are rabbit polyclonal sera produced during this study while Anti-Str mAb (**Right**) is a commercial anti-Str mouse monoclonal antibody. The green color indicates fluorescence of the HRP substrate while red indicates the prelabelled protein standards. Lanes: **A**, Ova; **B**, Ova-Str-Eda conjugate; **C**, Ova-Str-Eda conjugate; **D**, Lys; **E**, Lys-Str-Eda conjugate; **F**, Lys-Str-Eda conjugate.

#### Str-Csa

The ratio of Csa to Str (10:1 ratio) was predicted to form more dimeric adduct as long as the disulfide bond in Csa reoxidized with another monomeric Csa derivative. The LC profile revealed an elution peak that was composed primarily of mass species corresponding to reduced Str (584.29 Da as a +1 ion)(JM-0), and dimeric (Str)_2_ -Csa (1284.6 Da as the +2 ion)(JM-6); monomeric reduced Str-Csa (718.32 Da as a +1 ion)(MM3) was also detected (**Fig. 7, bottom**). While we assumed that borohydride reduction would cleave the cystamine (Csa) linker, apparently, oxidation during dialysis did drive disulfide bond formation producing a predominant peak of the dimeric JM-6 product and much less of the monomeric JM-3 product.

We found that the mass species detected as products in the three synthetic reaction samples tracked accurately with the predicted structural formulas and molecular weights of the expected chemical species (**Fig. 6 and 7**). Dihydro streptomycin or JM-0 (reduced with sodium borohydride) provided a comparator in this LCMS analysis. By measuring the abundance ratios of dimeric species to their corresponding monomeric modified species, we calculated that 61.3% and 38.3% of the Str-Csa preparation and Str-LinkB preparation were composed of dimers JM-6 and JM-4, respectively. In contrast, very little of the dimer JM-5 could be detected in the Str-Eda preparation. Comparing the ratios of predicted monomers to dihydro (reduced) Str in each preparation was also consistent with this conclusion with JM-1/JM-0 being 1.4 suggesting that JM-1 (the monomeric derivative with a free primary amino group) was likely the largest component of the Str-Eda reaction products. This conclusion was supported by the observation that the Str-Eda reaction products (after dialysis and lyophilization) reacted with di-nitro-benzene sulfonate (TNBS), a reagent that detects primary amino groups. Compared to unmodified Str which displays only a secondary amine, TNBS reacted much more strongly with components present in the Str-Eda preparation at pH 9.0. Accordingly, the Str-Eda reaction products that contained JM-1 were utilized for synthesis of protein conjugates by coupling its exposed primary amino group (**Fig. 6, Top**) to the episilon amino groups of lysine displayed by carrier proteins.

### Synthesis of OVA-Str-Eda and LYS-Str-Eda conjugate immunogens for generation of anti-Str hapten polyclonal antibodies

We used glutaraldeyde crosslinking to make a Str-protein conjugates between Str derivatives present in the Str-Eda reaction product mix and two carrier proteins hen egg white lysozyme (LYS) and ovalbumin (OVA) (see METHODS). The LYS-Str-Eda and OVA-Str-Eda conjugates we made were provided to a commercial vendor (Thermo Fischer Laboratories) who prepared polyclonal rabbit sera that were used in our experiments below. These included Day 1 (pre-immune sera) as well as hyper-immune, multi-boosted Day 72 sera.

In order to confirm that the anti-Str polyclonal sera provided by our commercial vendor detected Str as a hapten, we performed a western blot analysis on our Str-protein conjugates that included our vaccine immunogens. Anti-sera raised against LYS-Str-Eda or OVA-Str-Eda were used to probe membranes displaying the separated components of un-crosslinked carrier proteins as well as the protein-Str conjugates. Proteins were detected by silver staining of gels (not shown) while antibodies were detected with appropriate anti-rabbit IgG secondary antibodies coupled to horse radish peroxidase. As shown in **Fig. 8**, anti-OVA-Str-Eda sera detected a ‘ladder’ of higher molecular weight electrophoretic species in the LYS-Str-Eda conjugate preparation (corresponding to dimers, trimers, and larger crosslinked oligomers) but critically not monomeric LYS. Similarly, anti-LYS-Str-Eda sera detected a ladder of crosslinked species in the OVA-Str-Eda conjugate preparation but not monomeric LYS. Analysis using a commercial anti-Str mouse monoclonal antibody also confirmed that LYS-Str-Eda and OVA-Str-Eda conjugates were displaying the Str hapten as predicted. We conclude from these data, that Str was a good hapten in our conjugate immunogens and that these two rabbit antisera were suitable to evaluate the ability of polyclonal anti-Str antibodies to bind to cells and to promote complement-dependent killing of bacteria exposed to dimeric Str derivatives such as those present in the Str-Csa and Str-LinkB preparations.

### Str HARMs and anti-Str antibodies elicited by Str-carrier protein conjugate immunogens display bacteriocidal activity in the presence of human complement

To determine if anti-Str conjugate vaccine sera could kill bacterial cells in the presence of human complement, we utilized rabbit pre-immune (Day 0) sera as controls and compared it to fully boosted immune sera provided by the vendor (Day 72) who used our OVA-Str-Eda and LYS-Str-Eda conjugates as immunogens. *E. coli* E11Δpga StrR was used as a target in these studies because this strain is resistant to >32 μg/ml of Str and products of the the Str-derivative reactions performed (Str-Eda, Str-LinkB, and Str-Csa). We found that both Str-Csa and Str-LinkB reaction products could trigger modest (∼10-fold) killing of E11Δpga StrR in the presence of either anti-LYS-Str-Eda or anti-OVA-Str-Eda sera (**Fig. 9**). Str-Eda while more active than unmodified Str, was not as robust as Str-Csa and Str-LinkB in these antibody-dependent killing assays. We note that the dimeric HARM molecules in our Str-Csa and Str-LinkB preparations were not purified for these preliminary proof of concept experiments and may have even been antagonized by monomeric derivatives in the tested samples. Thus, these results, in principle, support the HARM concept for design of antibody recruiting molecules.

**Fig. 9.**
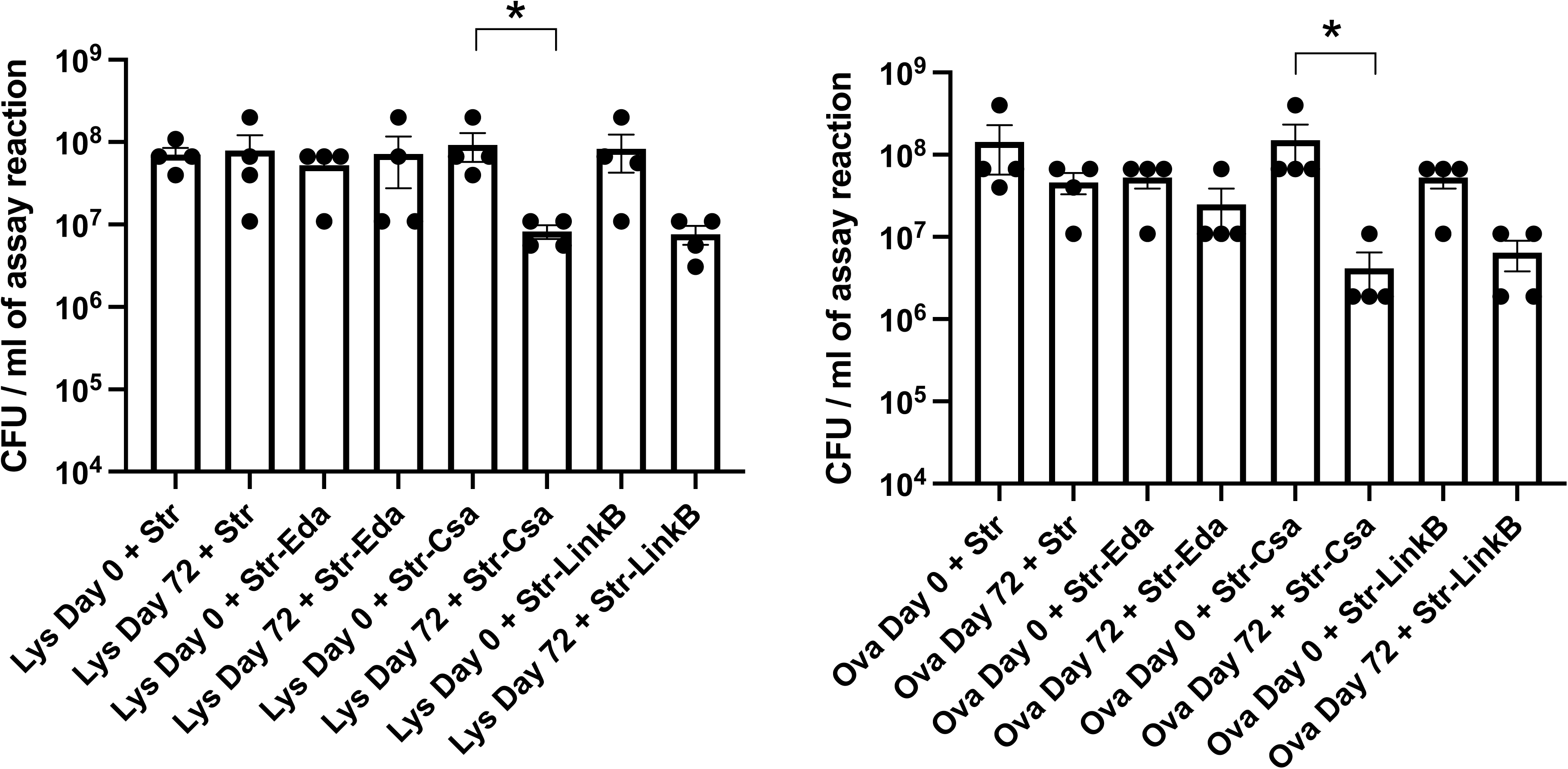
Bactericidal activity against *E. coli* in the presence of Str, derivatives of Str, and anti-sera against conjugate immunogens. Assays were performed on E11Δpga StrR in PBS at 37°C for 1 hr before viable counts were determined as CFU/ml remaining in the reaction. **Left**. Bactericidal activity of indicated combinations of Str and Str derivatives (Str-Eda, Str-Csa, and Str-LinkB) together with anti-OVA-Str-Eda conjugate rabbit sera collected at Day 0 or Day 72 of the immunization protocol (OVA labels X axis). **Right.** Bactericidal activity is shown of indicated combinations of Str, Str-Eda, Str-Csa, and Str-LinkB together with anti-LYS-Str-Eda conjugate rabbit sera collected at Day 0 or Day 72 of the immunization protocol (LYS labels X axis). Str and derivatives of Str were in the assay at a final concentration 2 μg/ml in the presence of the indicated rabbit antisera (final concentration 1%) and human complement (final concentration 0.5%). * Mann-Whitney test, value < 0.05 for indicated comparison data sets (brackets).

### Complement and anti-Str antibody killing of *P. aeruginosa* in the presence of dimeric Str HARM molecules

We also tested if anti-Str hapten antibodies and the HARM molecules present in the Str-Csa and Str-LinkB preparations could kill a complement sensitive strain of *P. aeruginosa* given that such strains represent a substantial percentage of clinical isolates particularly from pulmonary sites (19). Indeed, screening a subset of clinical isolates of *P. aeruginosa* showed that nearly 50% of these were sensitive to 25% human complement (**Fig. S1**). We therefore selected BWH9 for intial testing, a complement-sensitive, streptomycin-resistant isolate that was associated with a ventilator acquired pneumonia (VAP) infection (20). We found that components in the Str-LinkB and Str-Csa preparations (but not the Str-Eda preparation or unmodified Str) triggered significant killing of BWH9 in the presence of anti-LYS-Str-Eda or anti-OVA-Str-Eda rabbit antisera and 0.5% human complement (**Fig. 10**). The abundance of dimers JM-4 and JM-6 in Str-LinkB and Str-Csa preparations, respectively suggested they could explain the differences in potency observed.

**Fig 10.**
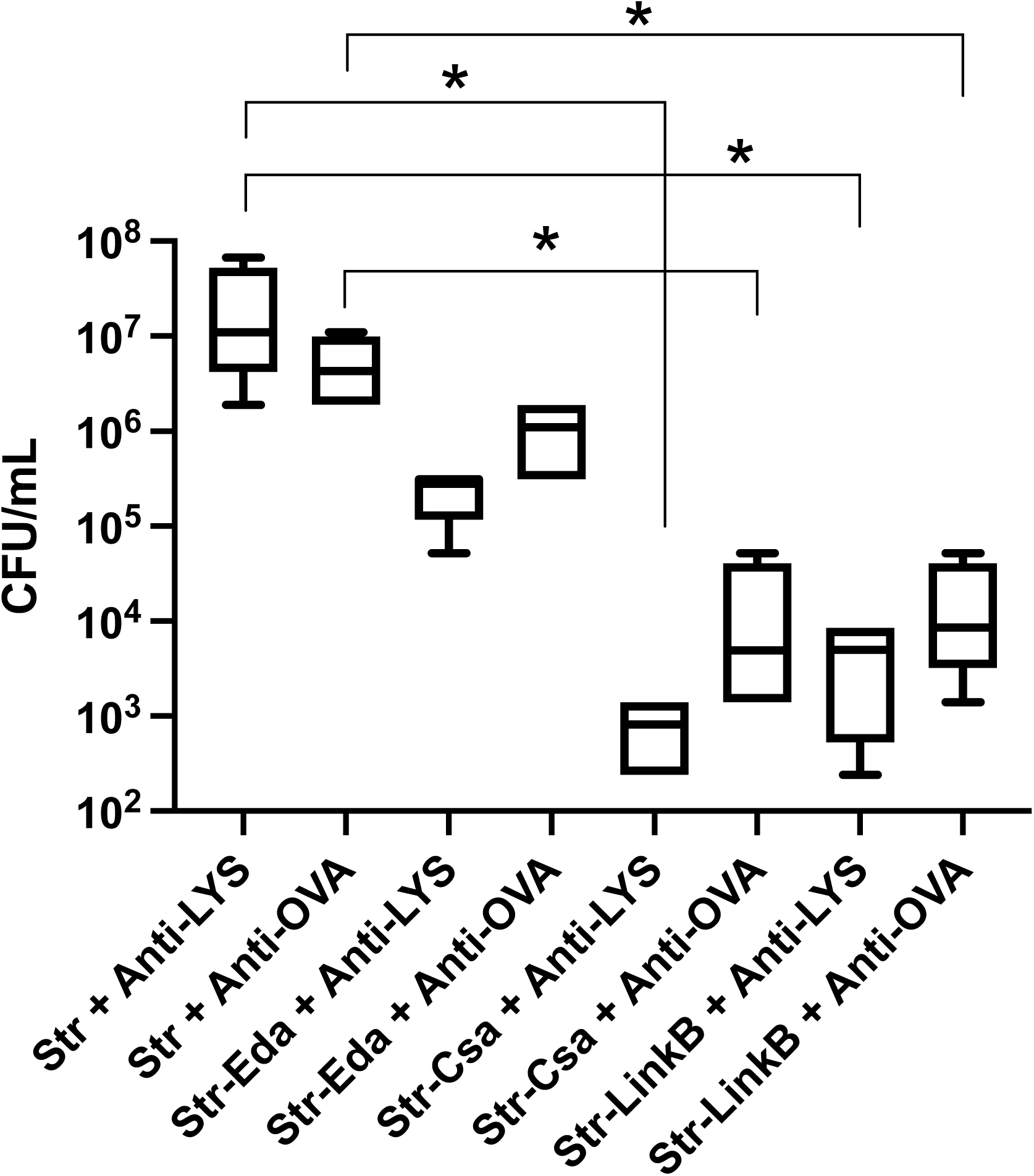
Sensitivity of *P. aeruginosa* to streptomycin derivatives and antisera against Str-Eda conjugate immunogens. Assays were performed on *P. aeruginosa* BWH9 in PBS at 37°C for 1 hr before viable counts were determined as CFU/ml remaining in the reaction. Bactericidal activity is shown of the indicated combinations of Str, and Str derivatives (Str-Eda, Str-Csa, and Str-LinkB) together with anti-LYS-Str-Eda conjugate sera (Anti-Lys) or anti-OVA-Str-Eda conjugate sera (Anti-Ova) collected at Day 72 of the immunization protocol. Str and derivatives of Str were in the assay at a final concentration 2 μg/ml in the presence of the indicated rabbit antisera (final concentration 1%) and human complement (final concentration 0.5%). * Mann-Whitney test, value < 0.05 for indicated comparison data sets (brackets).

Because the ARM preparation Str-Csa and polyclonal sera anti-LYS-Str-Eda were the two most potent components we identified in this initial investigation, we elected to study these two for enhancement of killing of BWH9 by human complement. As shown in **Fig. 11**, Str-Csa alone could enhance killing of BWH9 in the presence of human complement. The combination of Str-Csa and anti-LYS-Str-Eda antibody further enhanced killing by approximately 10-fold across several concentrations of human complement in the assays. We also found that reduction of Str-Csa with dithiothreitol eliminated its activity in the presence and absence of Anti-LYS-Str-Eda antibody, suggesting that the dimeric, disufide bonded form of Str-Csa was likely responsible for its enhanced bacteriocidal activity in these assays (data not shown). Together these results suggest that the dimeric form of Str-Csa (**JM-6, Fig. 6**) can enhance complement-dependent killing of *P. aeruginosa* BWH9 and that this bacteriocidal effect is enhanced by the presence of anti-Str hapten antibodies.

**Fig. 11.**
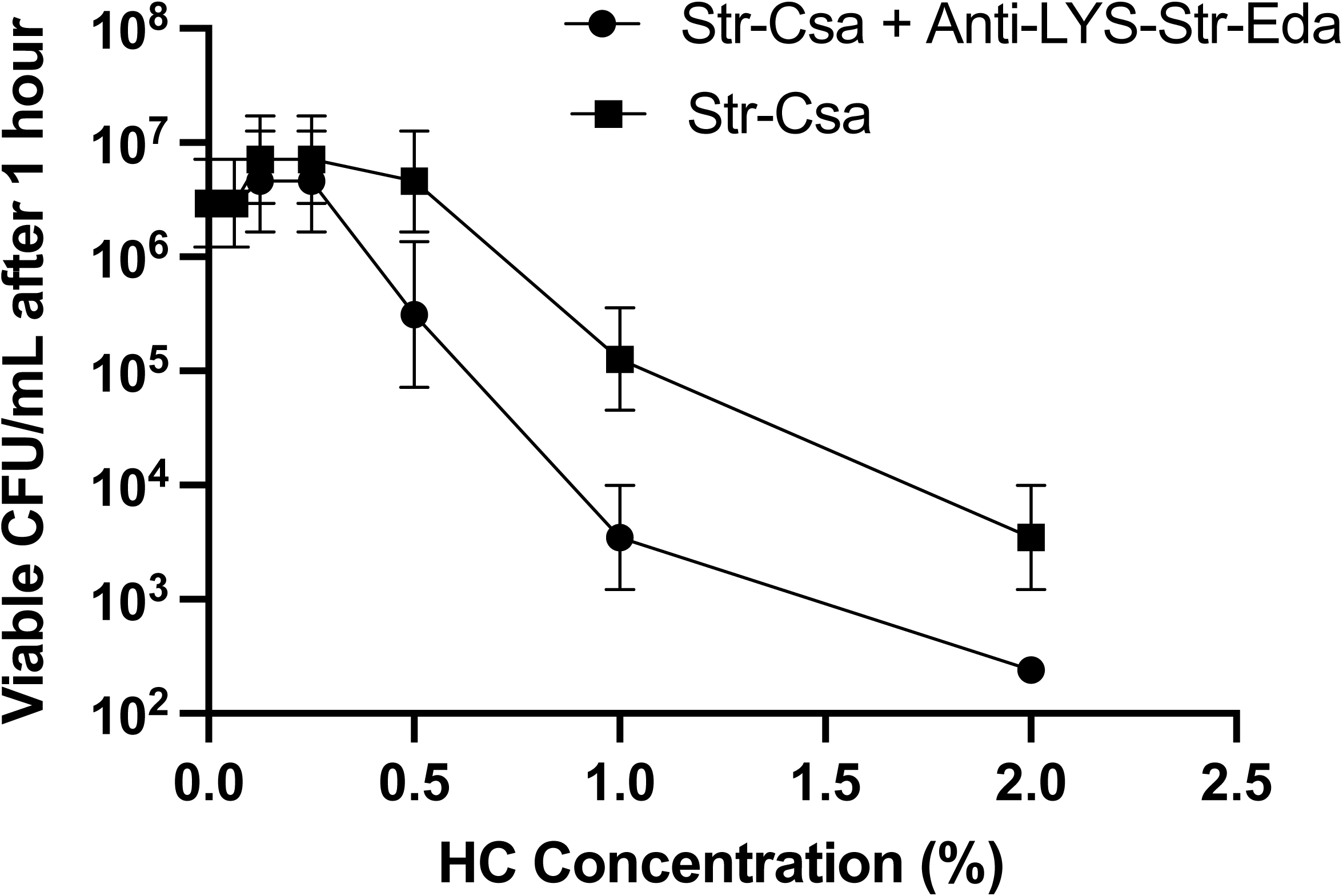
Bacteriocidal activity of human complement for *P. aeruginosa* as a function of the presence of Str-Csa or Anti-LYS-Str-Eda plus Str-Csa. The assays were performed in quadruplet on *P. aeruginosa* BWH9 in PBS at a starting concentration of 5 x 10^6^ per ml with either Str-Csa alone (2 μg/ml final concentration) or Str-Csa mixed with 1% anti-LYS Str-Eda conjugate sera (Day 72 bleed) at the indicated concentrations of human complement (HC). After 1 hr at 37°C, the assays were serial diluted to determine remaining viable bacteria per ml of the reaction mix.

## DISCUSSION

The data presented here provide further evidence that the ARM (antibody recruiting molecule) concept (6–13) shows promise as an approach to address the growing problem of MDR Gram-negative infections. One class of ARM molecules described here are synthetic peptides designed to display the A12X mimotope (16) of the O1 antigen of *V. cholerae*, a polysaccharide that is a major immunogenic component of the live attenuated vaccine PanChol (17) and likely other cholera vaccines as well (35). The cationic K6 peptide used to target this O1 mimotope ARM molecule to the bacterial cell surface is a antimicrobial peptide (AMP) that likely binds to the anionic Lipid A moeity on the surface of Gram-negative cells as either a monomer or in the form of nanomicellar structures (14). Local administration of K6 is active in treating infections due to *P.aeruginosa* (14) but the role of the host complement system in this *in vivo* activity of K6 has not been studied.

We found that both K6 and FZ5 (the ARM peptide of K6 fused to the A12X mimotope of O1) could sensitive some strains of Gram-negative cells to killing by human complement in the presence and absence of anti-O1 antibody. To observe this we had to use a laboratory strain of *E. coli* (OmniMAX) or a derivative of the *E. coli* K1 encapsulated clinical isolate E11 defective in production of poly-β-(1–6)-N-acetyl-D-glucosamine (PNAG) (26). Both of these strains were found to be highly sensitive to human complement. Unfortunately, *E. coli* and *P. aeruginosa* strains that were highly resistant to human complement did not show enhanced killing by FZ5 that was dependent on complement and anti-O1 antibodies *in vitro* (data not shown). These observations complicate the interpretation of our data showing modest enhanced killing of two complement-sensitive strains of *E. coli* by the combinations of FZ5, human complement and either rabbit anti-O1 antibodies, or human sera from individuals vaccinated withPanChol (**Figs. 1-4**). We acknowledge that the potentiation of killing is a preliminary result that could be further tested through mutagenesis of the FZ5 sequence and identification of active human anti-O1 monoclonal antibodies that react with FZ5. However, even with this level of validation, FZ5 enhancement of complement mediated killing in the absence of anti-O1 antibody remains a complicating factor in interpretation of our results.

Polymyxin B (PMB) has been reported to sensitize Gram-negative cells to killing by the complement system present in mammalian serum (36). Thus, ARM molecules based on conjugation of epitopes to PMB (12) should be evaluated for their potential effects on complement killing independent of anti-epitope antibodies. Most published studies of bacterial targeting ARM molecules have not progressed to animal efficacy studies (37). It remains to be determined if FZ5 is active as a treatment in infection models dependent of anti-O1 antibodies and complement. We also plan to explore if C3 knockout mice or cobra venum treated mice (which depletes complement components C3 and C5) can provide evidence that complement is indeed necessary for any therapeutic potency of an ARM molecule such as FZ5 in the presence of anti-O1 antibodies. It is also worth noting that we considered substituting the *bona fide* O1 oligosaccharide minimal epitope (38) in place of the mimotope displayed by FZ5. However, coupling of a suitable synthetic O1 oligosaccharide to the K6 peptide would require difficult carboydrate chemistry that was beyond our team’s collective experience.

We propose that synthetic peptide chemistry is far more practical in exploring the mimotope targeting ARM concept. Many widely used human vaccines (e.g., covid19, flu, diphtheria, tetanus, and polio vaccines) have the potential to elicit antibodies that react with linear peptide epitopes derived from the protein components of these vaccine immunogens (39–44). Such linear peptides, which are recognized by polyclonal responses elicited by these licensed vaccines, could be future components to evaluate in ARM designs that utilize fusion peptides to bind to prokaryotic (e.g., pathogenic or commensal bacteria) or even eukaryotic target cells (e.g., neoplasm) to trigger antibody and complement dependent cytotoxicity of these cells by epitope or mimotope display.

We also developed ARM molecules based on homodimers of the antibiotic streptomycin (Str). We coined the acronymn ‘HARM’ (<u>H</u>omodimeric <u>A</u>ntibody <u>R</u>ecruiting <u>M</u>olecule) to emphasize their symmetrical structure compared to other ARM molecules described in the literature that are heterodimers in their design and functional attributes (7). While dimers of aminoglycosides have previously been reported (33), these molecules were designed to maintain their inherent antibiotic mechanism of ribosomal inhibition. Our preparations Str-LinkB and Str-Csa contained reduced streptomycin (ML**-0**), the monomers JM-2 and JM-3 but also substantial amounts of two dimeric HARMs based on mass spectroscopic analysis (JM-4 and JM6) (**Fig. 6 and 7**). *E. coli* E11Δpga StrR and *P. aeruginosa* BWH9 were found to be sensitive to products in the Str-LinkB and Str-Csa preparations in the presence of low levels of human complement (**Figs. 10 and 11**) but we did not prove that the dimers of Str present in the preparations were responsible for the observed bactericidal activity. This sensitization to complement killing could be enhanced by antibody that was raised to recognize the Str hapten on these HARM derivatives but only about 10-fold compared to complement alone. We speculate that the highly cationic nature of JM-4 and JM-6 is more important to their ability to activate human complement killing of target Gram-negative cells than their abiltiy to activate anti-Str epitope antibody-dependent bacteriocidal activity. Mechanistically, this may be analogous to the ability of cationic peptides such as polymyxins to trigger enhanced complement killing (36).

Str is a cationic molecule and dimers of Str are predicted to be +6 in net charge at physiological pH. We propose that cationic HARMs are capable of activating human complement killing of Gram-negative cells by disruption of their outer membrane (OM) in a way that promotes multiple killing mechanisms (**Fig. 12**). For example, this OM disruption could result in exposure of lipid A and LPS core sugars which could activate complement killing by lectin-dependent and alternative (i.e., antibody-independent) pathways, or by faciltitating the translocation of the membrane attack complex (MAC) of the complement system into deeper targets in the bacterial cell envelope such as its inner membrane. MAC translocation into the bacterial inner membrane is thought to be the rate limiting step in the measurable loss of viability in complement-dependent bacteriocidal assays that do not depend on bacterial opsonization and killing by phagocytic cells (45). Thus, HARM molecules based on cationic antibiotics could potentially be mutifunctional. They could kill cells directly by outer and inner membrane disruption, expose alternative complement-dependent pathway ligands, and also enhance the bacteriocidal activity of antibodies that recognize HARM epitopes bound to cells through host cell-independent (i.e., classical and alternative complement activation) and host cell-dependent (i.e., opsonization and ADCC) pathways. If the HARM molecules themselves have their own antibacterial activity independent of the above, then this constellation of bacteriocidal mechanisms may not be easy to defeat by common MDR resistance mechanisms such as efflux, drug degradation and target mutation (4,5).

**Fig. 12.**
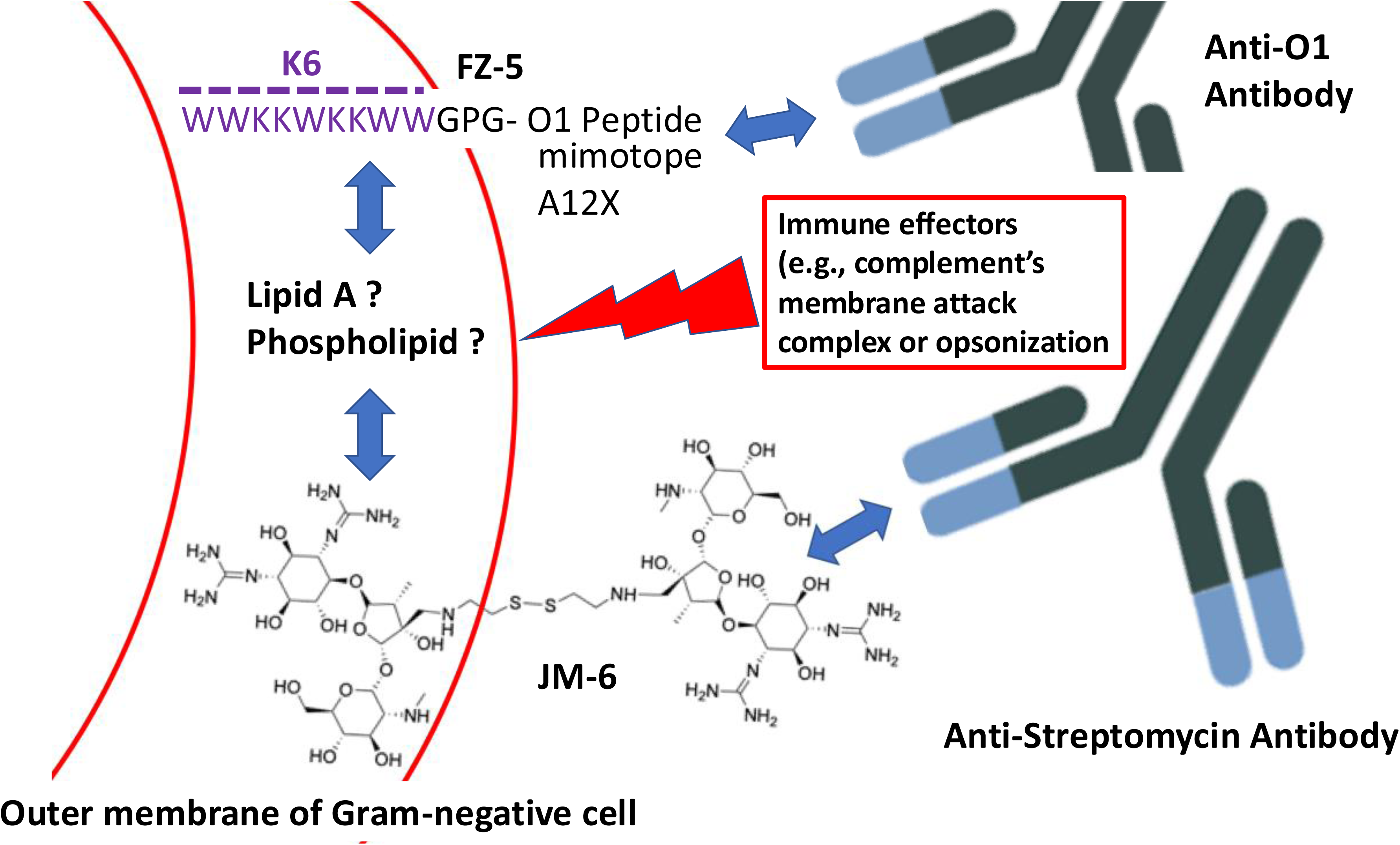
Schematic diagram showing hypothetical interactions of the ARM Peptide FZ-5 and HARM JM-6 with a Gram-negative bacterial cell, anti-O1, and anti-Str antibodies. The fusion peptide FZ-5 is proposed to bind to Gram-negative cells through its K6 moiety and then display its O1 mimotope (A12X) in ways that allows binding interactions (blue arrows) with anti-O1 antibodies. Similarly, the highly cationic, homodimeric, HARM derivative JM-6 is proposed to bind through one of its ends to anionic structures such as lipid A moiety of lipopolysaccharide or phospholipids in the outer leaflet of the Gram-negative bacterial outer membrane through ionic interactions. The distal portion of JM-6 acts as a haptin to bind to anti-Str antibodies recruiting them to the bacterial cell surface. If antibodies are recruited at sufficient local concentration, effectors of the immune system such as complement can attack the bacterial surface (red lightning bolt) by insertion of its membrane attack complex into the cell envelope or by triggering opsonization and killing of the bacterial through phagocytosis.

We also recognize that cationic antibiotics such as aminoglycosides and polymyxins have a checkered past because of their association with adverse events in treated patients (46). At this time it is unknown whether larger cationic molecules such as the dimeric Str derivatives described here will be safer or more toxic, or if anti-streptomycin antibodies would ameliorate or enhance adverse events associated with aminoglycosides. These same cautionary comments apply to ARM molecules based on the cationic peptide K6 described here or polymyxins (12). Serum sickness (allergic events associated with antibody-antigen complexes forming in the bloodstream) is another concern associated with the ARM approach (7). Maximum complement activation typically requires recruitment of the regulatory protein properdin after the initiation of the classical pathway near sites of multimeric antigen-antibody complexes (45, 50). However, it is likely that monomeric or dimeric hapten antibody interactions will not trigger enough antibody clustering in the bloodstream to allow classical pathway activation and subsequent properdin amplification during ARM-based therapies. Recruitment of antimicrobial ARM molecules to bacterial surfaces may also render their fluid phase concentration very low and thus dampen these theoretical adverse events associated with off-target complement activation in serum. Nonetheless, the field should remain mindful of these considerations if ARM therapeutics advance to clinical trials in the future.

In conclusion, our study provides new approaches to explore and extend the ARM concept. These include application of ARM molecules produced by simple peptide synthetic chemistry as well as antibiotic-derived, homodimeric ARM molecules (HARMs) that combine cell surface recognition with antibody recruitment. We hope these novel concepts will trigger more investigation into the use of ARM and HARM approaches in combating MDR bacterial infections as well as other diseases (e.g., cancer and autoimmunity) that would benefit from eliminating an offending cell type using the innate and adaptive immune systems.

## METHODS

### Strains and growth conditions

All bacterial strains used in this study are listed in the key resource table. Unless noted otherwise, bacteria were grown aerobically with 300 rpm shaking at 37°C in Luria-Bertani (LB) broth or on LB plates. Overnight cultures (ONC) were typically grown for 18 hr diluted 100-fold and grown for 5-6 hr more at 37°C before use in bacteriocidal assays as mid-log cells (OD_600_ of 0.2-0.4).

### Bacterial growth curve assays, complement and synthetic peptide sensitivity

Growth curves were generated by inoculating 96-well microtiter plates containing LB medium typically starting with an OD_600_ of 0.01. Cells were grown with continuous shaking at These solutions were adjusted to pH 9.5 and then cooled to 4°C. for 10 hours, with standard OD_600_ measurements taken every 10 mins using a BioTek Synergy H1 Plate Reader. When complement or peptide sensitivity was being assays, these were added before inoculation at indicated concentrations. Pooled human complement serum (Cat.#TCI-D1571) was obtained from Innovative Research. Peptides were obtained from GeneScript, Inc. and were processed for removal of trifluoroacetic acid by the supplier. Lyophilized peptides were dissolved in water before use.

### Chemical synthesis of streptomycin derivatives

Streptomycin (Str) sufate (Sigma) and other chemicals used in reactions were obtained from Sigma unless otherwise noted. Str was dissolved in 0.1M sodium borate buffer pH 9.5 at 100 mg/ml and then mixed with the three selected diamines at the following ratios: 1) Ethylene diamine (Eda) at 50-fold molar excess to Str, 2) Diethylene glycol bis (3-amino propyl) ether (LinkB) (Spectrum Chemical) at 20-fold molar excess to Str, and 3) Cystamine (Csa) at 10-fold molar excess to Str. The single unique aldehyde group displayed by Str was used as the target for a reductive amidation reaction in the presence of these three diamines (**Fig. 5**). Each solution of Str mixed with a given diamines was adjusted to pH 9.0 with dilute hydrochloric acid. Solid sodium borohydride was added to 50-fold molar excess to Str in each of the three reactions over the course of 30 mins at room temperature. After overnight incubation at 4°C, sodium sulfate was added to a final concentration of 10mM and the soluble fraction of each reaction was transferred to centrifuge tubes and adjusted to be 90% ethanol to precipitate Str-related derivatives. After 30 mins at -20°C, centrifugation at 5000g for 20 mins produced white pellets that quickly dissolved in water. The soluble products from each reaction were then separately dialyzed exhaustively against distilled water for 48 hr at 4°C using 500-1000 Da cutoff dialysis bags (Spectra Float-A-Lyzer G2 Dialysis Devise). Through control experiments, we determined that these dialysis membranes allowed passage of the three diamines used in our synthetic strategy but not unmodified Str. This apparently indicates that, like most carbohydrates, Str has a hydrodynamic volume in excess of its formula molecular weight. Tri-nitro-benzene sulfonate (TNBS) was used to test the contents that remained in the three bags after dialysis for the presence of primary amino groups. Whereas unmodified or sodium borohydride reduced Str did not react with TNBS, the products of the three diamine reductive amidation reactions were strongly positive after dialysis as expected for monomeric derivatives (which should display primary amino groups associated with the linkers used (**Fig. 6; JM-1, JM-2, and JM-3**). The contents of the bags were lyophilized and produced white powders that were stored at 4°C and are referred to in the text as Str-Eda, Str-LinkB, and Str-Csa reaction products.

### Liquid chromatography-mass spectrometry of Str derivatives

Str derivatives present in Str-Eda, Str-LinkB and Str-Csa reaction products were analyzed by liquid chromatography-tandem mass spectrometry (LC-MS-MS). In brief, 100 ng of sample was injected into a Vanquish UHPLC system (Thermo Fisher Scientific) equipped with an Eclipse Plus C18 column (4.6 × 75 mm, 1.8-µm particle size; Agilent Technologies) and coupled to a Q Exactive Orbitrap mass spectrometer (Thermo Fisher Scientific). The mobile phase A consisted of water containing 0.1% formic acid, and mobile phase B consisted of acetonitrile containing 0.05% formic acid. The gradient was programmed as follows: 1% B from 0 to 2 min, increased to 10% B at 7 min and 60% B at 9 min, held at 60% B until 10 min, returned to 1% B at 10.5 min, and held at 1% B until 15 min. The mass spectrometer was operated in positive-ion mode. Full-scan mass spectra were acquired over an m/z range of 200–2,500 at a resolution of 70,000, with an automatic gain control (AGC) target of 3 × 10⁶ and a maximum injection time of 100 ms. Tandem mass spectra were acquired at a resolution of 17,500, with an AGC target of 1 × 10⁵, a maximum injection time of 100 ms, an isolation window of 0.5 m/z, and a lower scan limit of m/z 50.

### Synthesis of conjugate vaccines composed of Str-Eda attached to carrier proteins and production of anti-Str haptin rabbit antisera

LC-MS-MS analysis indicated that the Str-Eda reaction contained the highest fraction of monomeric diamine-Str ligation product (corresponding to JM-1 in **Fig. 6**). Accordingly, this sample was used to prepare Str-protein conjugates vaccines. Ovalbumin (OVA), hen egg white lysozyme (LYS) were separately dissolved at 20 mg/ml each in 0.1M sodium acetate buffer, pH 5.0. Each of these protein solutions received an aliquot of Str-Eda stock to give a final concentration 9 mg/ml of reaction products in each solution. Glutaldehyde (5%) aliquots were added over a 30 min period at room temperature until the final concentration was 1% glutaraldhyde in each reaction. During this period the clear protein solutions became visibly opalescent, indicative of molecular crosslinking of OVA and LYS monomeric proteins and therefore presumably also attachment of the ethylene amine displayed by Str-Eda product JM-1 to the epsilon amino side chains of random lysine residues on OVA and LYS. Crosslinking was terminated after by the addition of glycine to 0.1M and incubation overnight at 4°C. The reactions were then dialyzed in 10,000 Dalton cutoff membrane bags overnight at 4°C against phosphate buffered saline (PBS). Approximately 10mg of the Str-Eda-LYS and Str-Eda-OVA conjugates was then delivered to a commercial vendor (Thermo Fischer Laboratories) to prepare polyclonal rabbit sera that were used in our experiments and referred to as anti-LYS-Str-Eda or anti-OVA-Str-Eda sera in the manuscript. These sera were heat inactivated at 56°C for 30 mins to remove complement activity in bactericidal assays.

#### Pooled human PanChol antisera sera

Human sera from volunteers who received one dose of the experimental vaccine PanChol were collected on Day 1 or Day 15. The geometric mean fold rise of the 5 participants at day 15 was 1552 (95% CI 582-4141). The immunization protocol previously approved by the Mass General Brigham Healthcare Institutional Review Board (2022P000461) and Institutional Biosafety Committee (2022B000015) and conducted under a US Food and Drug Administration (FDA) Investigational New Drug Application (28217) and is described in Leitner et al., (17). The two serum pools, Day 1 and Day 15 were blinded to the donors and maintained at -80°C until the day of use in bactericidal assays.

### Bactericidal assays for complement, peptides, or Str-derivatives in the presence or absence of antisera

Minimal inhibitory concentrations (MIC) assays were typically performed in LB broth where a component being tested (peptide or human complement were typically serially diluted 2-fold and then approximately 10^5^ bacteria were added to each test well. Typically, each assay was done in biological triplicates. After incubation at 37°C for 18 h, endpoints of growth were recorded based on optical density and the average concentration of drug or complement that eliminated growth was recorded as the MIC.

Assay reactions designed to detect rapid killing of bacterial cells by peptides or Str-derivatives in the presence or absence of complement and antisera were performed in 100 μl l volumes of Hank’s phosphate buffered saline in 96 microtiter plate wells. Components were typically assembled into wells in the following order: 1) PBS, 2) compounds being tested (e.g., Str-derivatives), 3) anti-sera (e.g., anti-O1 *V. cholerae* sera (Difco), Str-Eda-protein conjugate sera, or pooled human sera from PanChol vaccine recipients), 4) bacterial cells, and 5) human complement. All components were added from appropriate stocks to give the indicated final concentrations in either μg/ml for Str-derivatives, dilution factor of anti-sera, or % human complement. Fully assembled assays were incubated with 200 rpm rotational agitation at 37°C for 60 mins. Reactions were then diluted in LB broth to determine viable cell count (reported as colony forming units per ml of reaction) as determined by the endpoints of growth after 1:6 or 1:3 serial dilutions. Each assay was typically performed in 4 biological or technical replicates. Data obtained were plotted and analyzed for statistical significance using appropriate analytics available in PRISM software.

### Western Blot

Samples were prepared in 2X Laemmli buffer (Millipore Sigma) and boiled at 95°C for 10 minutes. Following cooling on ice, proteins in samples were separated by SDS-PAGE and transferred to a nitrocellulose membrane (iBlot2, Thermo Fisher Scientific). Membranes were blotted with primary (e.g., anti-Str-Eda protein conjugates or anti-Str mouse monoclonal GTX39076 antibody (Genetex). Fluorescent secondary antibodies were used (Licor IRDye 680RD and IRDye 800CW) and blots were imaged using a Licor Odyssey Imager. ImageStudio v4.0.21 was used for data analysis.

## AUTHOR CONTRIBUTIONS

Conceptualization: JJM

Methodology: FGZ, JJM, MLM

Investigation: FGZ, JJM, MLM, DRL, SRW

Visualization: FGZ, JJM, MLM

Funding acquisition: JJM, MKW

Project administration: JJM, MKW

Supervision: JJM, MKW

Writing – original draft: JJM

Writing – review & editing: FGZ, MLM, MKW, SRW, DRL, JJM

## ACKNOWLEDGEMENTS

This work was supported by National Institute of Allergy and Infectious Diseases RO1 Grant AI-018045 (to J.J.M.) and the Howard Hughes Medical Institute (HHMI) (to M.K.W.). This article is subject to HHMI’s Open Access to Publications policy. HHMI lab heads have previously granted a non-exclusive CC BY 4.0 license to the public and a sublicensable license to HHMI in their research articles. Pursuant to those licenses, the author-accepted manuscript of this article can be made freely available under a CC BY 4.0 license immediately upon publication. This manuscript is the result of funding in part by the National Institutes of Health (NIH). It is subject to the NIH Public Access Policy. Through acceptance of this federal funding, NIH has been given the right to make this manuscript publicly available in PubMed Central upon the Official Date of Publication, as defined by NIH. The authors sincerely thank Dr. Tiandi Yang for his generous assistance in LCMS analysis and Dr. Wen Zheng for her training of M.L.M. in use of the graphic and statistical software packages Prism, Chemdraw, and Powerpoint. We also thank Dr. Gerald Pier for sharing *E. coli* strain E11 and its Δpga mutant for our studies.

## Supplemental Figures

**Fig. S1.**
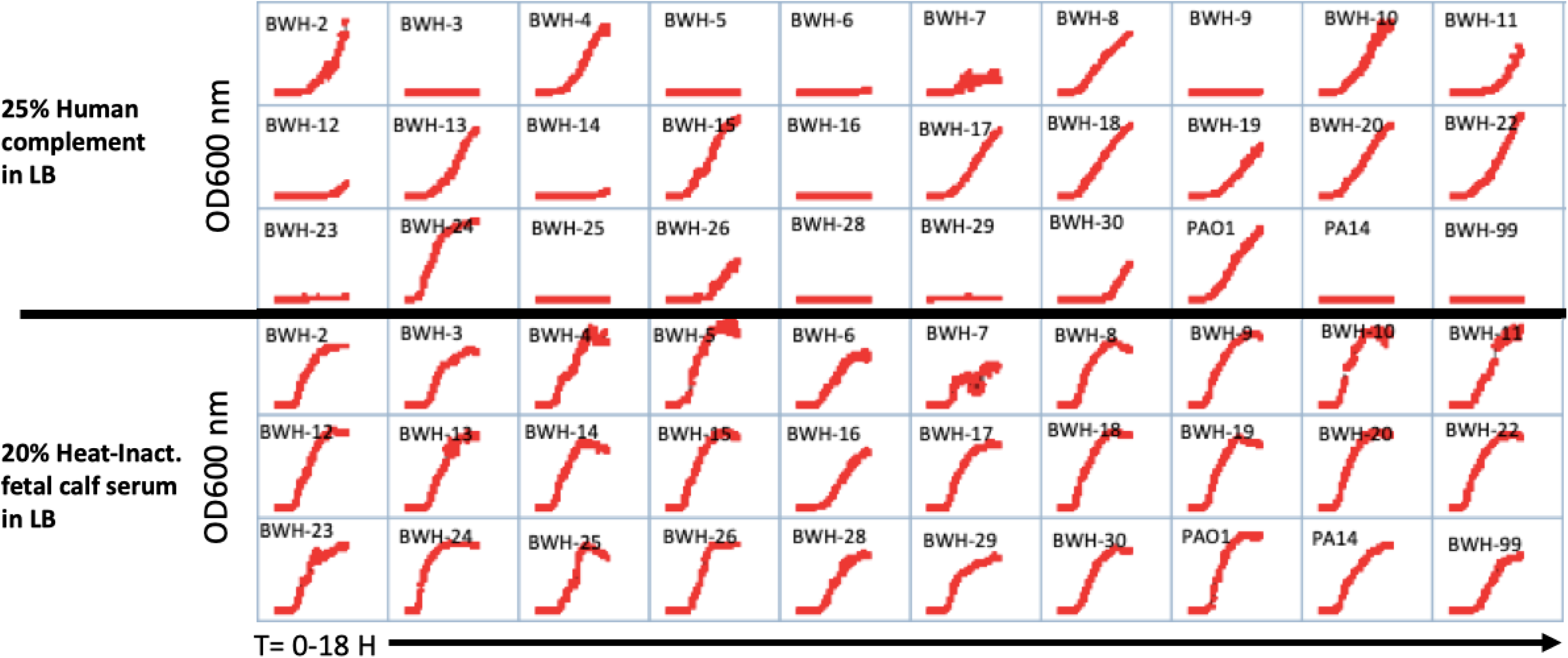
Sensitivity of clinical isolates of *P. aeruginosa* to human complement. Growth curves (OD600 measurements over 18 hr at 37°C) of the indicated isolates of *P. aeruginosa* obtained from Brigham and Women’s Hospital (BWH) (20) is shown in the presence of 25% final volume human complement obtained from Innovative Research, Inc. (top) or in the presence of 20% final volume heat-inactivated fetal calf serum in Luria broth (LB) (bottom).

